# “Visualize, Explore, and Select”: A Protein Language Model-based Approach Enabling Navigation of Protein Sequence Space for Enzyme Discovery and Mining

**DOI:** 10.64898/2026.03.23.712833

**Authors:** Felix Moorhoff, David Medina-Ortiz, Alicja Kotnis, Ahmed Hassanin, Mehdi D. Davari

## Abstract

The rapid expansion of protein sequence databases continues to outpace functional characterization, creating a persistent bottleneck in enzyme discovery and mining—particularly in large, heterogeneous, and sparsely annotated sequence spaces. This gap is amplified by visualization challenges and the lack of informed strategies for exploration, selection, and mining across sequence spaces. Here, we present an embedding-based workflow, implemented in a computational platform called SelectZyme, for alignment-free visualization and exploration of protein sequence space that combines protein language model (pLM) representations with dimensionality reduction and hierarchical density-based clustering. The approach links complementary visualizations of protein sequence space as low-dimensional landscapes, connectivity projections (minimum spanning trees), and dendrogram-based organization, enabling coherent interactive exploration and candidate selection across global context and local neighborhoods without relying on sequence-identity thresholds, EC numbers, conserved motifs, or predefined functional annotations. Across distinct case studies, we demonstrate that embedding-defined neighborhoods remain structurally conserved even when sequence identity falls within the ‘twilight zone’, and that coherent functional organization emerges also for modular protein segments in a fully unsupervised analysis. We also show how this workflow supports user friendly, interactive and scalable enzyme mining in a sparsely annotated, complex multi-family protein space surpassing *>* 100,000 sequences, enabling constrained candidate selection around experimentally validated anchors. By enabling interactive exploration across visualizations, supporting informed candidate selection, our workflow streamlines biocatalyst discovery and helps to bridge uncharacterized sequence-space to functional characterization campaigns - thus providing a broad starting point to downstream protein engineering, machine-learning–guided design cycles, and iterative experimental screening campaigns.

## 1 Introduction

Enzymes are indispensable catalysts in biotechnology, enabling essential transformations in medicine, energy conversion, environmental remediation, and industrial manufacturing [Robinson, 2015]. Their large sequence and structural diversity, shaped by billions of years of evolution across heterogeneous ecological niches, provides a vast repertoire of biocatalysts spanning broad ranges of substrate specificity, temperature tolerance, pH optima, and stability profiles [Kim et al., 2025, Yeo et al., 2025, Durairaj et al., 2023, Yang et al., 2025, Barrio-Hernandez et al., 2023]. While this diversity represents a remarkable resource for biocatalysis, it also poses a fundamental challenge, since reliably identifying and prioritizing catalytically relevant enzymes within an ever-expanding and heterogeneous sequence landscape remains difficult [Moorhoff et al., 2025].

Despite the rapid growth of protein sequence databases, functional characterization continues to lag far behind sequence accumulation. The majority of available entries remain uncharacterized or rely on automated annotations with limited experimental validation [Bateman et al., 2024, Shinde et al., 2024]. This annotation gap constrains the practical exploitation of enzyme diversity, complicates the identification of promising biocatalysts, and limits the pool of starting points for downstream protein engineering campaigns aimed at tailoring industrially valuable biocatalysts [Zhou et al., 2024, Yang et al., 2024]. Enzyme mining and disocvery has therefore emerged as a central computational strategy to bridge this gap by prioritizing promising candidates from genomic, metagenomic, and specialized databases, particularly in scenarios where experimental labels are sparse or absent [Moorhoff et al., 2025].

Conventional enzyme mining workflows typically follow a sequence of steps that includes database retrieval, similarity-based characterization, functional inference, and experimental validation, as illustrated in Figure **1**A [Bateman et al., 2024, Sayers et al., 2024, Richardson et al., 2022]. However, at the scale and diversity of modern sequence repositories, the characterization stage presents substantial limitations. These include visualization bottlenecks, extensive manual parameter tuning, reliance on alignment-based similarity metrics, and the potential loss of local diversity through clustering and representative selection (Figure **1**B) [Hong et al., 2024, Hamamsy et al., 2023].

**Figure 1:**
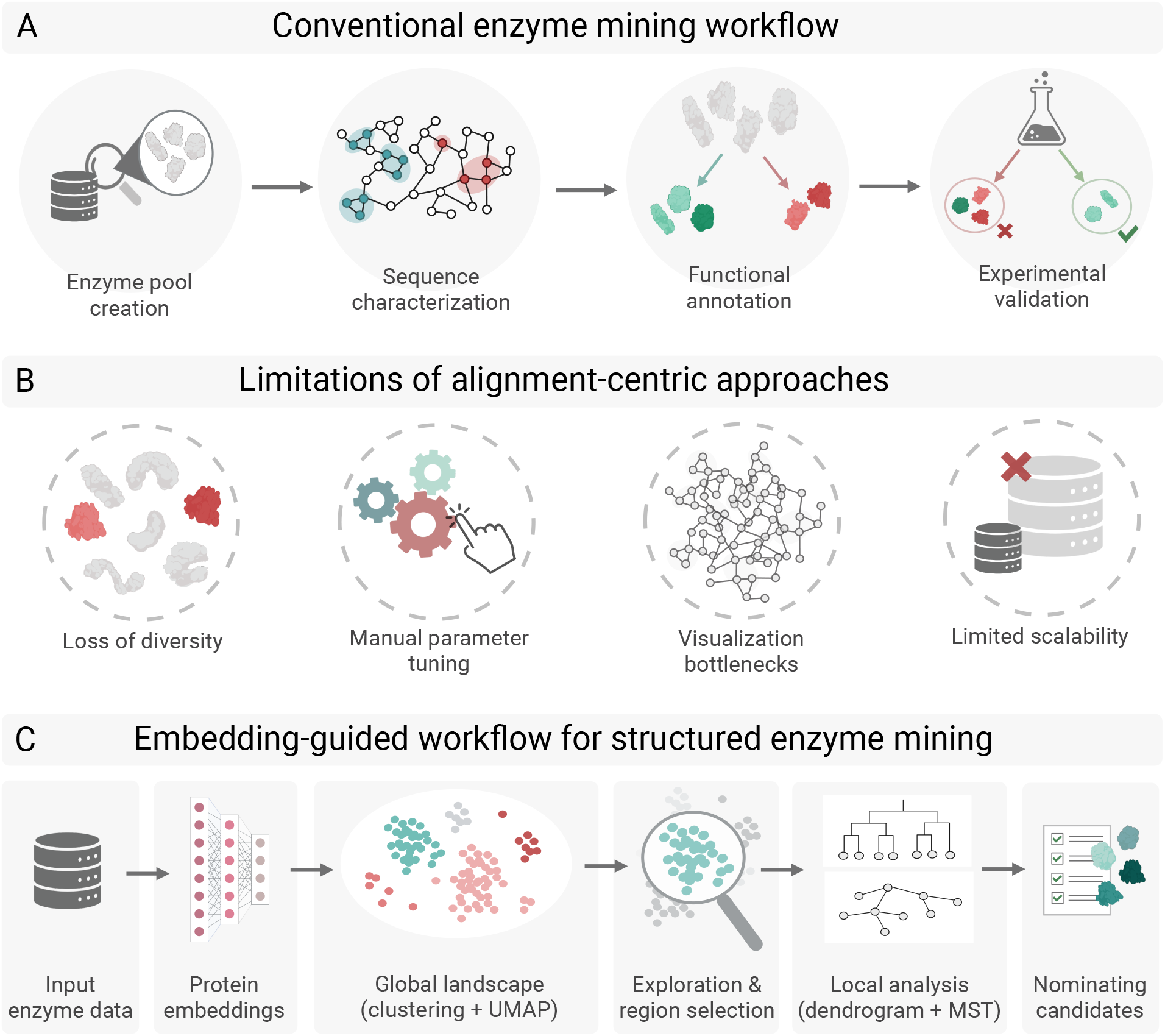
From alignment-centric enzyme mining to embedding-guided structured navigation. **A** Conventional enzyme mining pipelines typically proceed through sequential stages that include enzyme pool creation via database retrieval, sequence characterization through pairwise similarity or multiple sequence alignment, functional annotation based on conserved features or predictive models, and experimental validation using biochemical assays or high-throughput screening. **B** At large scale, alignment-centric workflows face inherent limitations, including loss of biological diversity through representative-based clustering, reliance on manual parameter tuning, visualization bottlenecks in dense similarity networks, and limited scalability as dataset size increases. **C** Embedding-guided workflow proposed in this study. Sequences are first embedded using protein language models to construct a global landscape via density-based clustering and dimensionality reduction. Exploration then proceeds through iterative region selection, connectivity restoration using minimum spanning trees, hierarchical interpretation via dendrograms, and structured candidate nomination. This formalized navigation framework enables scalable, hypothesis-driven enzyme mining while preserving both global context and local neighborhood coherence in large and heterogeneous sequence spaces.

Sequence comparison strategies based on sequence alignment, including local similarity search tools such as BLAST [Altschul et al., 1990] and multiple sequence alignment approaches, have long served as the foundation for protein comparison. Yet alignment-centric strategies have become increasingly impractical and less informative for large families and superfamilies, particularly under low sequence identity and variable sequence length [Hamamsy et al., 2023]. Redundancy reduction approaches based on predefined similarity thresholds, including CD-HIT Huang et al. [2010], MMseqs Mirdita et al. [2019], DBSCAN [Ester et al., 1996], and clustered reference resources such as UniRef Suzek et al. [2007] and RefSeq Pruitt [2004], O’Leary et al. [2015], Chen et al. [2016], improve scalability but may obscure functional diversity by collapsing heterogeneous neighborhoods into single representatives.

Sequence Similarity Networks (SSNs) extend pairwise similarity analysis by representing proteins as nodes connected through alignment-derived edges [Zallot et al., 2019, Copp et al., 2019]. While powerful in moderate-size datasets, SSNs often become dense and highly sensitive to threshold selection as dataset size increases, making them difficult to visualize and interpret while inheriting alignment-based constraints [Hornung and Terrapon, 2023, Chen et al., 2025]. As highlighted by Gerlt et al. [2015], networks containing several million edges can correspond to only a few thousand sequences and already require substantial computational resources, underscoring the practical limits of graph-based visualization in highly connected spaces [Chen et al., 2025]. Integrated platforms such as EnzymeMiner [Hon et al., 2020] combine retrieval, property prediction, and network export, yet they may require substantial prior knowledge when annotations are sparse [Hon et al., 2020]. Additional background on sequence comparison strategies and enzyme pool construction is provided in the Supporting Information (Sections S2–S3 with additional practical considerations summarized in Figure S1).

Recent advances in protein language models (pLM) provide an alternative perspective by mapping sequences to high-dimensional embeddings that capture functional and structural information directly from sequence context [Rives et al., 2021, Erckert and Rost, 2024, Lin et al., 2023, Heinzinger et al., 2024, Senoner et al., 2025a, Su et al., 2023, Elnaggar et al., 2022]. These representations enable scalable exploration of large sequence collections in a manner that is independent of explicit alignments. Nevertheless, embedding-based approaches are rarely formalized as practical layers within enzyme mining workflows and often lack the task-oriented structure required for biologically grounded interpretation and candidate prioritization [Chen et al., 2025, Senoner et al., 2025b]. This limitation is particularly relevant because enzyme discovery is typically driven by clearly defined objectives, including organismal origin, substrate scope, product selectivity, or process conditions, which are difficult to incorporate into rigid alignment-centric pipelines [Robinson et al., 2021]. In parallel, the increasing application of machine learning to enzymatic property prediction has highlighted the risk of selection-biased training sets and the need for representative yet strategically constrained sampling strategies that balance assay feasibility with multiple-criteria optimization objectives [Fahsbender et al., 2025, Bernett et al., 2024, Bushuiev et al., 2024, Kapoor and Narayanan, 2023].

To address these challenges, we introduce a computational methodology that formalizes enzyme mining as a navigation approach in embedding space. The approach integrates embeddings derived from protein language models with dimensionality reduction and density-based clustering to construct a global representation of sequence space, followed by iterative exploration, connectivity reconstruction, and hierarchical analysis (Figure **1** C). Rather than replacing alignment-based analysis, this framework operates as a decision-support layer that guides users from global landscape inspection to local neighborhood analysis and candidate nomination under sparse annotation.

The methodology is implemented in the platform SelectZyme to ensure reproducible execution and interactive exploration. However, the primary contribution is not the software interface itself, but the explicit formalization of an embedding-guided workflow that links global landscape construction, region selection, connectivity-aware contextualization, and hierarchical interpretation. This integration preserves both global context and local structural coherence while avoiding rigid similarity thresholds.

Importantly, the framework does not prescribe a single mining strategy or objective. Instead, it supports multiple exploration modes tailored to distinct discovery objectives, including novelty-oriented exploration, optimization around close homologs, and constraint-aware filtering based on desired properties. By combining density patterns, community structure, connectivity reconstruction, and hierarchical distances, the workflow enables structured ranking of subregions and principled candidate selection. Practical constraints such as organism preference, annotation status, or functional descriptors can be incorporated as flexible filters rather than hard requirements (Additional implementation details and selection strategies are described in the Supporting Information Section S1).

To demonstrate the utility of the workflow, we show that coherent functional organization can emerge in a fully unsupervised manner from embedding-based proximity, even at the level of modular protein domains. We further demonstrate that large and heterogeneous enzyme families can be navigated under sparse experimental labeling through anchor-guided prioritization, while connectivity- and hierarchy-aware analyses provide structural insights that support rational candidate selection in catalytic discovery workflows. These results position embedding-guided navigation not simply as a visualization tool, but as a scalable and methodologically explicit framework for hypothesis-driven enzyme mining in expansive and weakly annotated sequence landscapes.

## 2 Results and discussions

### 2.1 Structured embedding-guided workflow for enzyme mining

We formalize enzyme mining as navigation and exploration process in embedding space, as summarized in Figure **1**C. Starting from user-defined sequence sets, protein language model embeddings are used to construct a global representation of sequence space, which is then structured through density-based clustering and dimensionality reduction. In contrast to alignment-centric pipelines and sequence similarity networks that rely on pairwise comparisons and manually selected similarity thresholds [Hornung and Terrapon, 2023], this framework operates in a latent space in which relatedness is expressed as proximity in high dimensional space rather than discrete cutoff-based connectivity.

Exploration proceeds iteratively from global landscape inspection to local neighborhood analysis. Regions of interest can be defined based on proximity to experimentally validated anchors, existing annotations from databases, density patterns, or other metadata filters. Because nonlinear projections do not preserve full relational structure, connectivity is explicitly reconstructed through a minimum spanning tree derived from hierarchical clustering of high dimensional embedding space [Susmelj et al., 2023, Gower and Ross, 1969]. This minimum spanning tree integrates global and local relationships by linking sequences according to their intrinsic embedding distances, thereby restoring relational continuity across regions that may appear separated in reduced two-dimensional projections [Chaudhuri et al., 2014]. Unlike threshold-defined similarity networks which are highly sensitive to parameter selection, this connectivity layer restores disintegrated regions [Hornung and Terrapon, 2023, Probst and Reymond, 2020]. More detailed information about this concept can be found in Section S3 of the Supporting Information.

Hierarchical interpretation via dendrograms further quantifies distances between sequences and subgroups, enabling principled assessment of inter- and intra-community separation [Theys et al., 2019]. Landscape visualization, minimum spanning tree connectivity, and hierarchical quantification transform enzyme mining into a structured decision process rather than a simple similarity filtering approach.

Candidate selection is performed by integrating embedding proximity, cluster context, connectivity relationships, hierarchical distance, and optional metadata constraints. This multi-layered framework preserves both global context and local coherence while avoiding arbitrary similarity cutoffs.

By maintaining graded relatedness across distant sequence regimes and restoring relational structure, the workflow provides a scalable foundation for hypothesis-driven enzyme discovery in large and sparsely annotated sequence spaces. The following sections demonstrate how this framework enables unsupervised functional organization and supports prioritization under realistic discovery constraints.

### 2.2 Representative Applications of the Embedding-Guided Workflow

#### 2.2.1 Unsupervised emergence of functional organization in LOV sequence space

To evaluate whether the embedding-guided workflow can recover biologically meaningful organization in a fully unsupervised setting, we analyzed a curated dataset of proteins containing light–oxygen–voltage (LOV) domains. LOV domains constitute a good proof-of-principle system, as modular blue-light sensing elements distributed across diverse taxa and are coupled to multiple effector architectures [Glantz et al., 2016, Kopka et al., 2017]. Their architectural heterogeneity provides a stringent test of whether functional structure can emerge from embedding space without supervision nor use of explicit domain-level information [Kopka et al., 2017].

The dataset was derived from the curated LOV compendium reported by Glantz et al. [2016], comprising approximately 5,500 protein sequences annotated with distinct effector types and functional groupings. Importantly, no annotation labels were used during embedding generation, clustering, or dimensionality reduction. Furthermore, domain boundaries were not specified; the analysis operated exclusively on full-length protein embeddings. This setup ensures that any observed organization arises intrinsically from latent representation of full length sequences rather than supervised grouping or manual feature engineering.

Quantitative assessment of functional grouping using k-Nearest-Neighbor (kNN) agreement reveals strong local coherence for functional descriptors (Figure **2**A). Primary Effector, and Functional Cluster annotations exhibit substantially higher neighborhood consistency than expected under label randomization (baseline). In contrast, fine grained taxonomic descriptors such as Class and Order display weaker coherence, while unique accession identifiers behave as negative controls with expected minimal agreement. These results indicate that the embedding space is able to capture substantial differences of unseen properties and preferentially organizes sequences according to functional properties as discriminators rather than taxonomic relatedness or arbitrary identifiers.

**Figure 2:**
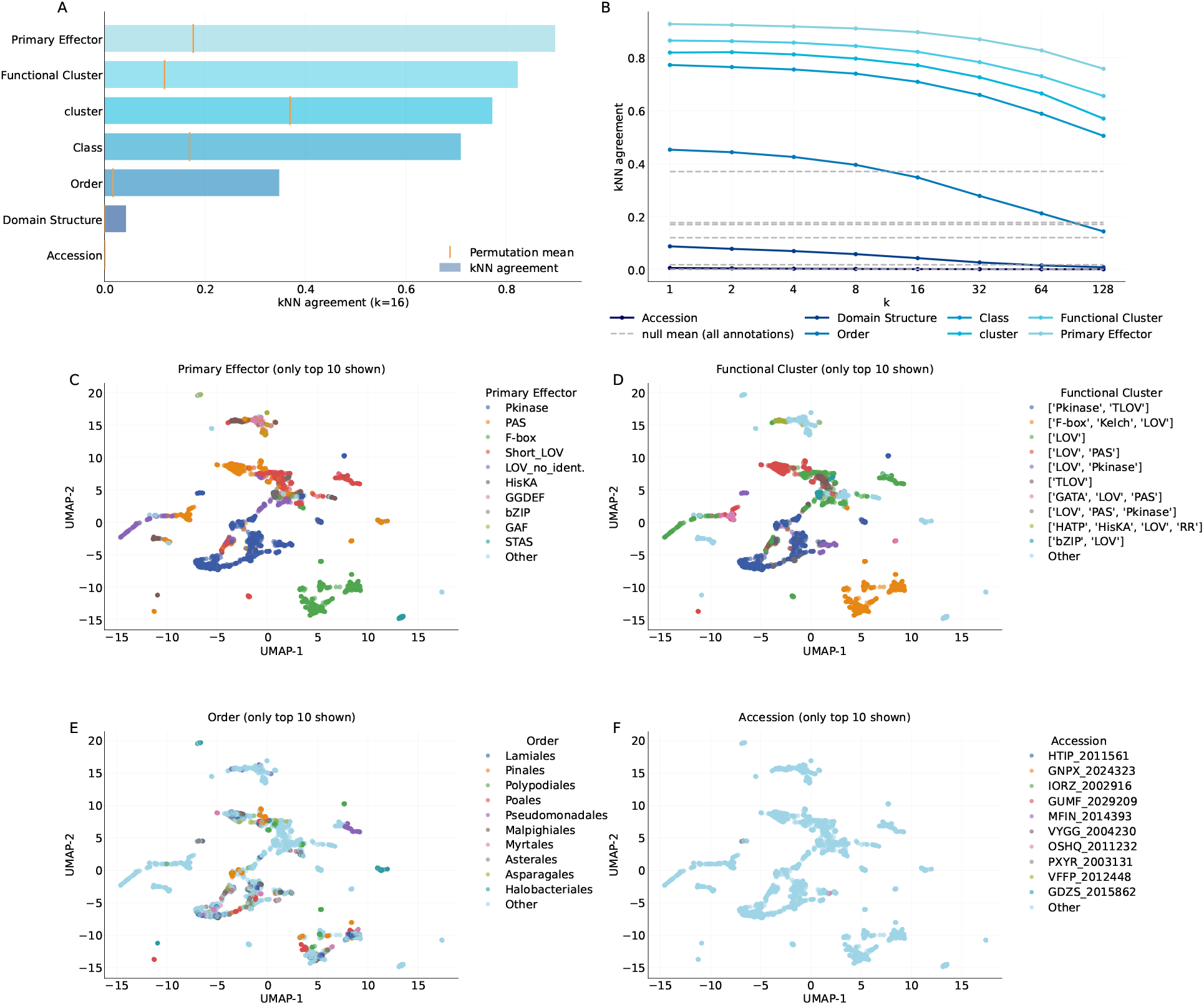
Unsupervised functional organization in LOV-domain proteins. **A** kNN agreement at *k* = 16 as a measure of local label consistency for selected LOV annotations, shown together with permutation mean. Functional descriptors (Primary Effector, Functional Cluster) display substantially higher neighborhood coherence than selected taxonomic descriptors or unique identifiers. **B** Robustness analysis of kNN agreement across varying neighborhood sizes (*k* = 1–128). Functional annotations consistently exhibit stronger neighborhood agreement than taxonomic descriptors across a wide range of *k*, indicating that the observed organization is not driven by a specific hyperparameter choice. Given the dataset size (approximately 5,500 sequences), large neighborhoods such as *k* = 128 already aggregate relatively broad sequence communities, which can gradually reduce agreement by merging smaller functional groups. Lines connect the evaluated *k* values for visualization purposes and do not represent interpolated measurements. Permutation-based null expectations remain independent of *k*, reflecting solely the class bias. **C–F** UMAP projections colored by representative annotations, illustrating strong (C,D) and weak (E,F) neighborhood coherence consistent with quantitative kNN agreement results. For readability, only the 10 most abundant classes are shown in the plots, while remaining labels are grouped as *Other*. All UMAP projections share the same coordinate system.

To assess robustness, kNN agreement was evaluated across a broad range of neighborhood sizes (Figure **2**B). Functional annotations consistently maintain higher agreement than taxonomic descriptors from very local neighborhoods (*k* = 1) to substantially broader community scales (*k* = 128). Given the total dataset size of roughly 5,500 sequences, a neighborhood size of *k* = 128 already represents a relatively large fraction of local neighboring sequences. At this scale, smaller functional clusters begin to merge, leading to a gradual reduction in agreement due to the aggregation of heterogeneous groups. The permutation-based null mean remain unaffected by *k*, as they are reflecting the original class bias for each annotation.

Visualization of the low-dimensional embedding further supports these findings (Figure **2**C–F). Annotations associated with effector type and functional cluster form coherent and spatially confined regions, whereas taxonomic labels exhibit diffuse and overlapping distributions. The agreement between quantitative neighborhood metrics and geometric organization patterns underscores that functional structure emerges intrinsically from embedding proximity.

This case study demonstrates that the proposed workflow can recover meaningful functional organization in a fully unsupervised setting, even for modular domains embedded within heterogeneous sequence contexts. The results validate the embedding-guided landscape as a biologically informative representation and establish a foundation for subsequent large-scale mining under sparse labeling conditions.

#### 2.2.2 Large-scale multi-family enzyme mining in PETase sequence space

Polyethylene terephthalate (PET) is one of the most extensively produced synthetic polymers and a major contributor to persistent plastic pollution, motivating enzymatic depolymerization as a sustainable alternative to conventional recycling strategies [Wei et al., 2025b]. PET-hydrolyzing activity is not confined to a single enzyme family but has been reported across multiple related hydrolase classes, most prominently cutinases, lipases, and carboxylesterases [Buchholz et al., 2022]. As a consequence, the PETase-relevant sequence space is intrinsically heterogeneous spanning multiple families and sparsely annotated, representing a challenging large-scale mining challenge [Seo et al., 2025, Turak et al., 2025].

To anchor exploration in experimentally validated evidence, we integrated curated literature data on plastic-degrading activity [Medina-Ortiz et al., 2025]. Enzymes experimentally confirmed to exhibit PET-hydrolyzing activity, as well as enzymes reported to lack such activity, were designated as *anchors*, while all remaining sequences were treated as unlabeled. This configuration reflects practical discovery conditions in which validated examples are limited and prioritization must rely on guided navigation of relational organization rather than dense annotation.

Protein language model embeddings were projected into a two-dimensional representation using UMAP to construct a global embedding landscape of the PETase-related sequence space via SelectZyme. The resulting space reveals a multi-family organization in which carboxylesterases, lipases, cutinases and esterases define distinct yet partially overlapping regions of sequence space (Figure **3**A). Overlaying curated activity annotations across the same embedding highlights localized neighborhoods enriched for PETase-positive and PETase-negative anchors (Figure **3**B), providing a reference for navigating the surrounding sequence landscape.

**Figure 3:**
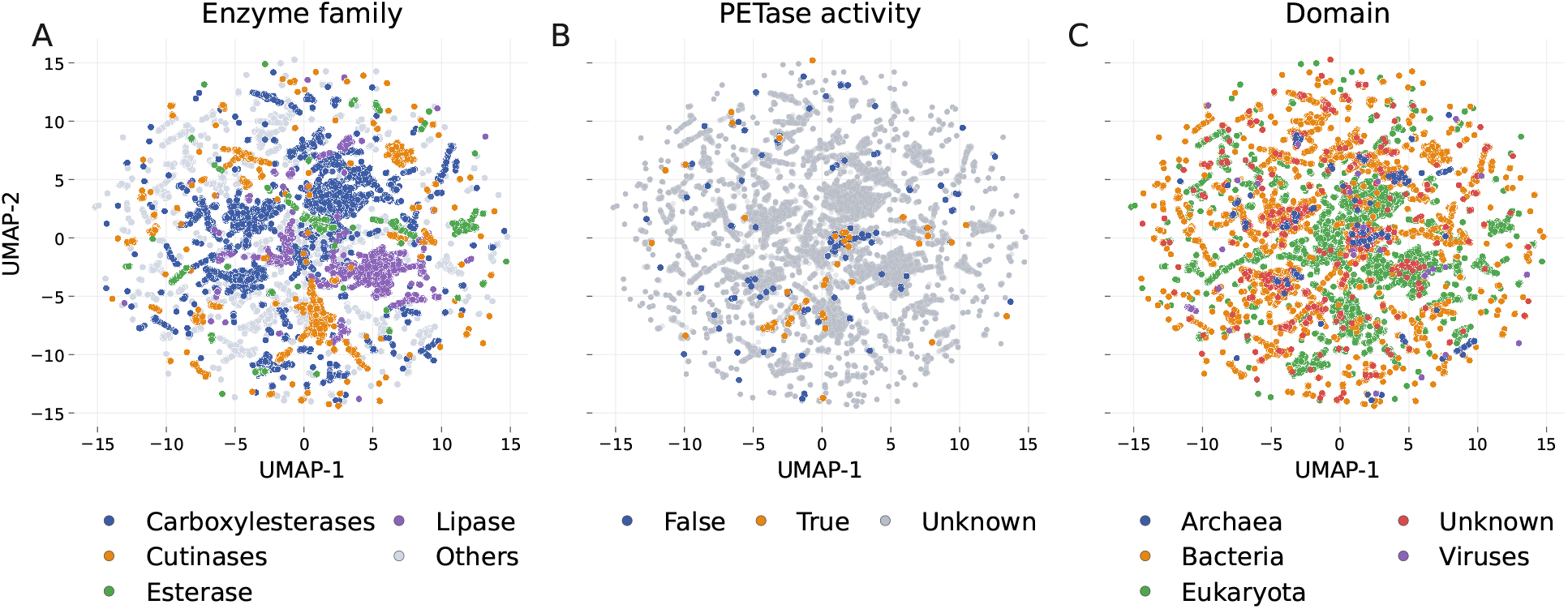
Embedding-guided navigation of heterogeneous PETase sequence space. **A** Two-dimensional visualization of the protein embedding landscape obtained by applying UMAP to protein language model embeddings of a multi-family dataset of PETase-related enzymes. The resulting landscape reveals a structured organization in which carboxylesterases, lipases, cutinases and esterases form distinct yet partially overlapping regions in sequence space, reflecting the broader functional diversity surrounding PET-degrading enzymes. **B** Overlay of curated PETase activity annotations across the same embedding landscape. Experimentally confirmed PET-active enzymes and reported non-active enzymes define localized anchor regions, while the majority of sequences remain unlabeled. This configuration highlights regions of sequence space enriched in PETase-positive neighborhoods that may contain additional candidate enzymes. **C** The embedding colored by taxonomic domain, enabling identification of archaeal sequences within PETase-proximal neighborhoods. Because archaeal proteins are frequently associated with increased thermostability and environmental robustness, these regions provide potential starting points for prioritizing candidate enzymes with improved biophysical properties. All UMAP projections share the same coordinate system.

Industrial PET depolymerization imposes stringent operational constraints, including elevated temperatures, progressive acidification during hydrolysis, and in some contexts increased salinity [Wei et al., 2025a]. Because such properties are rarely directly annotated, we adopted taxonomic origin as a pragmatic proxy for robustness. Exploration was therefore restricted to embedding-defined neighborhoods proximal to PETase-positive anchors, while prioritization favored *archaeal* sequences as candidates for enhanced thermostability and stress tolerance (Figure **3**C). This approach illustrates how the embedding-guided workflow supports enzyme mining under sparse and indirect functional constraints by combining anchor proximity with biologically motivated filtering criteria. Additional information on dataset construction and candidate selection criteria is provided in Supporting Information Section S4.

#### 2.2.3 Connectivity-aware refinement within PETase candidate neighborhoods

A refined subregion of the PETase landscape contains both PETase-positive and PETase-negative anchors, providing a controlled setting for assessing candidate prioritization beyond visual inspection of low-dimensional projections. While the embedding landscape captures global organization, two-dimensional projections alone do not fully preserve the relational structure of the underlying embedding space (Figure **4**A).

**Figure 4:**
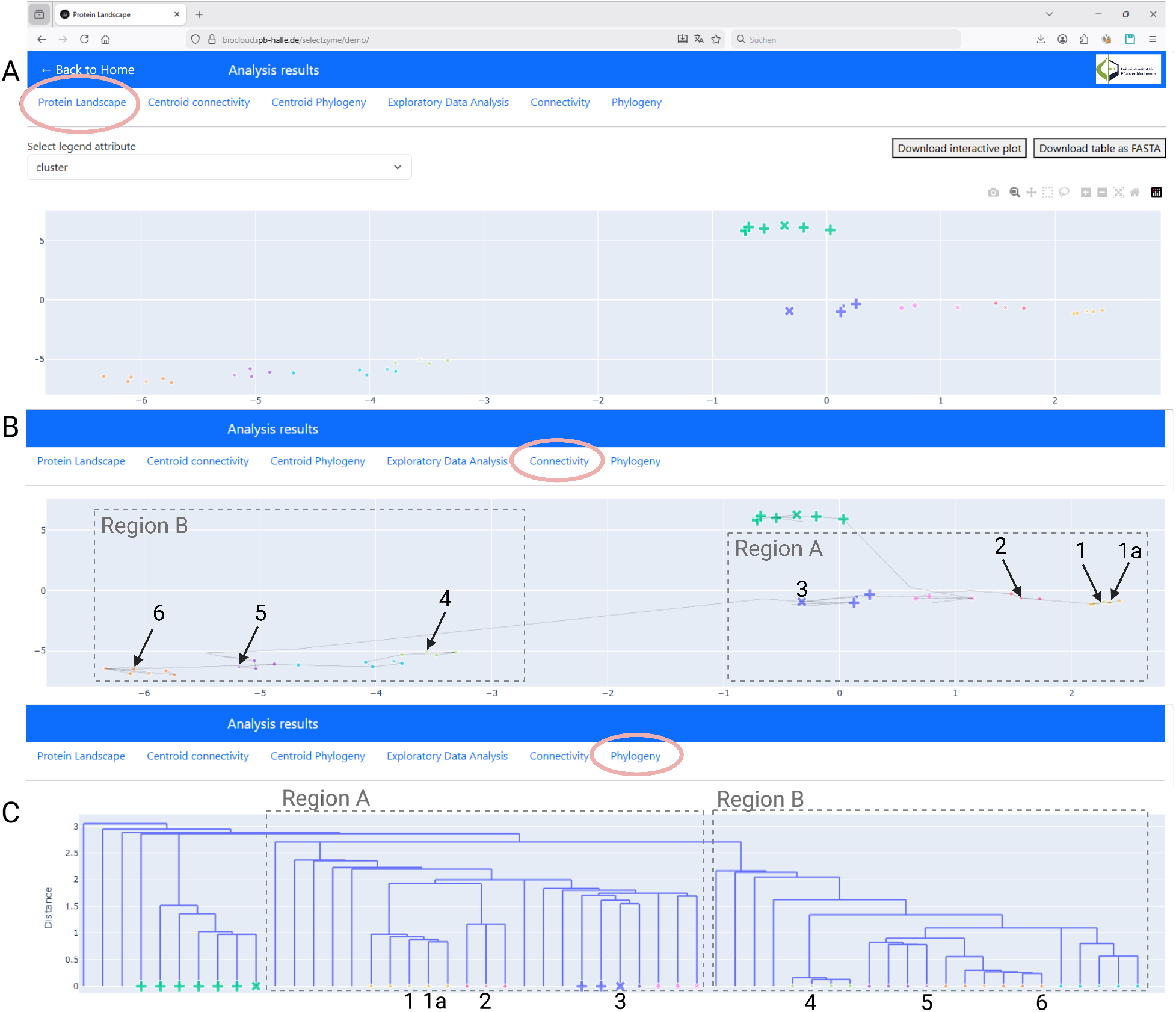
Connectivity-aware refinement of candidate prioritization in PETase sequence space. **A** Two-dimensional embedding landscape provides an intuitive global overview of protein sequence space but does not fully preserve hierarchical relationships or long-range connectivity. **B** Reconstruction of embedding-space connectivity using a minimum spanning tree derived from hierarchical clustering restores relational structure across the landscape without imposing similarity thresholds, linking locally coherent neighborhoods across distant regions. **C** Hierarchical organization of the same embedding space visualized as a dendrogram enables quantitative interpretation of subgroup structure and relative distances that are not apparent in two-dimensional projections. These combined complementary views illustrate how SelectZyme integrates embedding visualization to support candidate prioritization across distinct visualizations and objectives. For interactive exploration, an online version of this demonstration is available at https://biocloud.ipb-halle.de/selectzyme/demo/.

To refine interpretation, we reconstructed connectivity using a minimum spanning tree derived from hierarchical clustering in embedding space (Figure **4**B). This reconstruction restores graded relationships across locally coherent neighborhoods and clarifies how sequences are positioned relative to PETase-positive anchors within the embedding landscape.

Within this refined region, several candidates that appear proximal in the two-dimensional projection are resolved into distinct relational branches when hierarchical structure is considered. Conversely, sequences that appear visually separated remain connected through short embedding-space paths, indicating functional or evolutionary continuity not immediately evident in the projection. These distinctions affect prioritization by differentiating closely related variants from structurally or evolutionarily more distant neighbors.

Hierarchical distances further quantify separation between local communities (Figure **4**C). Short branch lengths correspond to tightly clustered variants, whereas longer hierarchical transitions mark shifts between embedding-defined subgroups. This quantitative layer enables discrimination across similarity regimes without imposing rigid identity cutoffs.

By integrating projection, connectivity reconstruction, and hierarchical distance, candidate selection becomes grounded in relational contexts rather than visual proximity alone. In heterogeneous multi-family settings such as PETase sequence space, this connectivity-aware refinement step reduces ambiguity in anchor-guided prioritization and mitigates the instability associated with threshold-based similarity networks. Importantly, these analyses are implemented within the SelectZyme framework, which provides both the computational workflow and an interactive visualization environment for exploring embedding landscapes and relational structure.

#### 2.2.4 Beyond sequence-level similarity

Embedding-based representations enable scalable navigation of protein sequence space. However, their value for enzyme discovery ultimately depends on whether proximity in embedding space reflects biologically meaningful relatedness. To assess this relationship, we compared embedding-derived neighborhood relationships with conventional sequence- and structure-based similarity metrics for a representative subset of sequences sampled from PETase-proximal regions of the embedding landscape.

Pairwise sequence comparisons derived from multiple sequence alignment (MSA) reveal limited residue conservation across the selected candidates (Figure **5**A). Although a tiny conserved core is detectable, the corresponding heatmap of pairwise sequence identities indicates that most sequence pairs fall within the low-identity regime (Figure **5**B). The majority of identities lies below 30%, placing many comparisons within the so-called twilight zone, where homology inference based solely on sequence identity becomes unreliable [Rost, 1999]. Only immediately adjacent variants display higher similarity, such as sequences 1 and 1a (63% identity) and sequences 5 and 6 (59% identity). Under a sequence-identity-based interpretation, several of these candidates could therefore be classified as weakly related or potentially belonging to distinct families.

**Figure 5:**
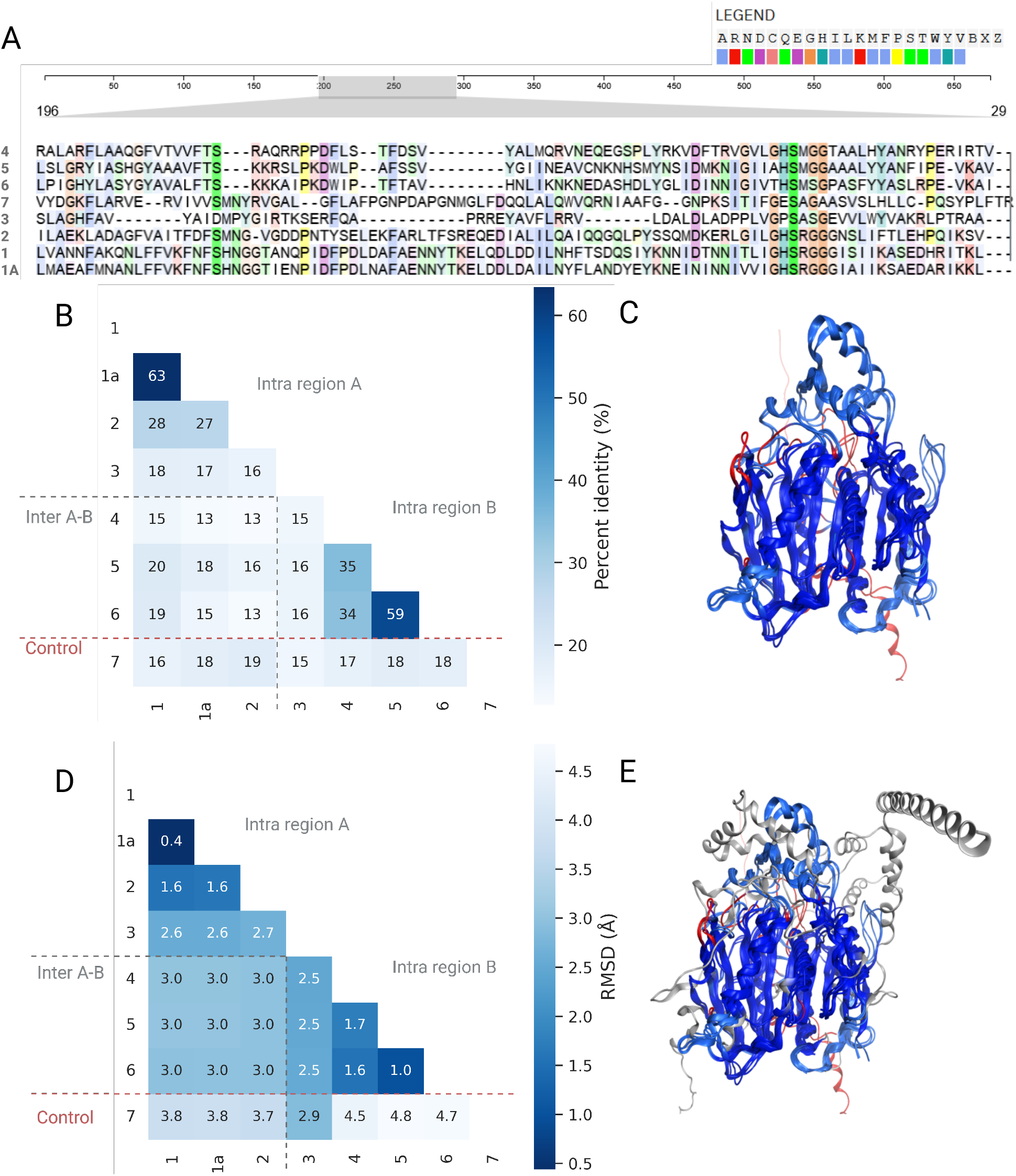
Embedding proximity reflects structural conservation beyond sequence identity. **A** Excerpt of the multiple sequence alignment (MSA) for representative PETase-proximal candidates, illustrating limited residue conservation across the selected sequences. **B** Heatmap of pairwise sequence identities derived from the MSA, showing that most comparisons fall within the low-identity regime (*<* 30%), corresponding to the twilight zone of sequence similarity. **C** Structural overlay of representative candidates demonstrating conservation of the core fold despite substantial sequence divergence. **D** Heatmap of pairwise structural distances obtained using TM-align, revealing gradual variation in RMSD across embedding-defined neighborhoods rather than abrupt structural divergence. **E** Structural alignment including a negative control sequence, which introduces additional non-alignable secondary structure elements (gray), hence confirms that the observed structural similarity among candidates is not shared across distant regions of the dataset.

In contrast, structural comparison reveals a conserved core architecture across representative candidates (Figure **5**C). The heatmap of pairwise structural distances obtained via TM-align shows gradual variation in root mean square deviation (RMSD) rather than abrupt structural divergence (Figure **5**D). For example, sequences 1 and 1a exhibit minimal structural deviation of approximately 0.4 Å and remain structurally similar to sequence 2 despite belonging to a different local subgroup. Comparable structural coherence is observed among sequences 4, 5, and 6, even in cases where sequence identity is comparatively low.

Inclusion of a negative control further clarifies this relationship. Structural alignment reveals additional non-alignable secondary structure elements in the control sequence (Figure **5**E), confirming that structural similarity is not trivially shared across unrelated proteins.

These observations indicate that embedding-derived proximity aligns more closely with fold-level structural conservation than with raw sequence identity in distant homology regimes. In other words, the learned representation appears to capture constraints associated with the underlying structural scaffold, even when primary sequence similarity falls within the twilight zone. This behavior is consistent with recent observations that protein language model embeddings encode information related to fold topology and higher-order structural features beyond residue-level conservation.

In heterogeneous enzyme families such as PETase-related hydrolases, where sequence similarity alone often becomes ambiguous, this representation-level signal provides a more stable basis for interpreting relatedness. Embedding-defined neighborhoods therefore serve as a structural proxy that complements traditional sequence comparison, enabling candidate prioritization within regions of sequence space where conventional similarity metrics provide limited guidance.

These results provide independent structural support for the biological relevance of embedding-defined neighborhoods. While the workflow leverages embedding-space organization to guide discovery across sparsely annotated enzyme families, its effective application still requires careful consideration of methodological choices, dataset composition, and domain-specific constraints, which we discuss in the following section.

### 2.3 Practical considerations, recommendations, and limitations

The analyses presented above illustrate how embedding-guided navigation can facilitate exploration of heterogeneous enzyme sequence spaces under sparse annotation. At the same time, the practical utility of the workflow depends on several methodological choices that influence robustness, and downstream prioritization.

First, the composition of the initial enzyme pool strongly constrains the landscape that can be explored. Embedding-based navigation cannot compensate for biased or incomplete sequence selection [Ding and Steinhardt, 2024]. Query design, database choice, and filtering criteria directly shape the embedding space and determine which neighborhoods become accessible for exploration [Ding and Steinhardt, 2024]. In practice, careful curation of the initial sequence pool is therefore critical, and iterative refinement of the input may be necessary to ensure representative coverage of diversity.

Second, the choice of protein language model can substantially influence the geometry of the resulting embedding landscape. Different pretrained models rely on distinct training corpora, architectures, and objectives, and may capture functional or structural signals with varying sensitivity [Bjerregaard et al., 2025, Ding and Steinhardt, 2024]. Comparative evaluation of alternative pLMs, model ensembles, or task-specific fine-tuning strategies may therefore improve robustness and domain adaptation [Senoner et al., 2025a]. However, systematic benchmarking across models represents a substantial effort in itself and is often better addressed as a dedicated methodological study rather than within a single enzyme mining application.

Third, embedding proximity should not be interpreted as a direct proxy for functional equivalence. Enzymatic activity ultimately depends on specific catalytic residues, substrate-binding pockets, oligomeric states, and environmental context, which may not be fully resolved at the level of global sequence embeddings [Flöge et al., 2025, Su et al., 2025]. Embedding-guided prioritization should therefore be viewed as a decision-support layer that complements, rather than replaces, structural modeling, biochemical validation, or experimental screening.

Finally, the workflow inherits limitations associated with dimensionality reduction and clustering. Although connectivity reconstruction and hierarchical analysis can mitigate projection artifacts, any two-dimensional visualization remains an abstraction of the underlying high-dimensional structure [Peng et al., 2025]. Interpretation of cluster boundaries or neighborhood transitions should therefore be approached with caution.

These considerations highlight that embedding-guided exploration is most effective when combined with careful dataset design, complementary analytical methods, and domain-specific biological knowledge. When applied within such a framework, the approach can provide a practical foundation for navigating complex sequence landscapes and identifying promising candidates for further investigation.

## 3 Conclusions and Perspectives

We introduced an embedding-guided workflow for navigating heterogeneous enzyme sequence spaces that integrates protein language model representations with dimensionality reduction, clustering, connectivity reconstruction, and hierarchical interpretation. Rather than relying on rigid similarity thresholds or single-objective unsupervised optimization, the framework emphasizes relations across embedding space and enables coherent navigation between global landscape context and local neighborhoods through complementary visualization and analysis layers.

Across distinct case studies, the workflow recovered biologically meaningful organization under different levels of annotation density. In the LOV domain analysis, functional coherence emerged in a fully unsupervised setting despite operating exclusively on full-length protein embeddings without explicit domain segmentation. In the multi-family PETase landscape, the framework supported anchor-guided prioritization within a heterogeneous and sparsely labeled sequence space, including the identification of extremophile-derived candidates under process-motivated constraints. Independent structural comparisons further demonstrated that embedding-defined proximity remains aligned with fold-level conservation even in sequence identity regimes where alignment-based homology inference becomes ambiguous.

These results position embedding-guided navigation not merely as a visualization strategy, but as a practical decision-support layer for enzyme discovery in large and sparsely annotated sequence landscapes. By combining latent representations with connectivity-aware and hierarchical interpretation, the workflow offers an interpretable and scalable alternative to traditional sequence similarity network approaches while preserving sensitivity across distant homology regimes.

Looking forward, several extensions may further expand the utility of this framework. Integration of predictive models for enzymatic activity, stability, substrate scope, or other physicochemical properties could enable multi-objective scoring. Coupling embedding-guided exploration with active learning, iterative experimental feedback, or structure-informed representations may further strengthen the connection between computational navigation and empirical validation.

## 4 Methods

### 4.1 Dataset Construction

The LOV dataset was derived from Supplementary Table 4 of Glantz et al. [2016], which provides curated annotations for proteins containing light–oxygen–voltage (LOV) domains. The non-redundant sequence collection was extracted and reformatted to include standardized accession and sequence fields compatible with the SelectZyme input schema. No additional preprocessing or filtering was applied. The resulting dataset comprises 5,500 sequences annotated across 27 functional and taxonomic descriptors and served as a benchmark for evaluating the unsupervised emergence of functional organization in embedding space.

The multi-family PETase dataset was constructed by integrating experimentally curated plastic-degrading enzymes from Medina-Ortiz et al. [2025] with entries retrieved from PlasticDB [Gambarini et al., 2022], PAZy [Buchholz et al., 2022], and UniProt. Literature reports indicate that PET-hydrolyzing activity has been observed across cutinases (EC 3.1.1.74), lipases (EC 3.1.1.3), and carboxylesterases (EC 3.1.1.1, EC 3.1.1.2, EC 3.1.1.101). Representative InterPro and Pfam signatures associated with these enzyme classes were therefore used to retrieve family-level sequence pools, including PF01083 (Cutinase), IPR013818 and PF00151 (Lipase), IPR003140 (Phospholipase/Carboxylesterase/Thioesterase), and cd00312 (Esterase/Lipase). UniProt sequences were collected with a length constraint of 70–800 amino acids to exclude truncated fragments and unusually large multidomain proteins unlikely to represent canonical hydrolases. Overlapping motif hits were deduplicated to retain unique sequence entries. The curated benchmark set was incorporated as labeled anchor sequences. The final integrated dataset comprised 105,299 non-redundant sequences spanning multiple hydrolase families.

For local refinement analysis, an embedding-defined PETase-proximal subregion was delineated based on spatial proximity to experimentally validated anchors and archaeal candidates in embedding space. Sequences within this region were exported and reanalyzed independently using identical embedding and clustering procedures to preserve methodological consistency. A representative subset of seven sequences sampled from this refined neighborhood was selected for downstream sequence- and structure-level comparison.

### 4.2 Embedding Extraction

Protein sequence embeddings were generated using pretrained protein language models (pLMs). For the LOV dataset, embeddings were extracted using ESM-2 (650M parameters), while the PETase datasets were processed using ESM-1b (650M parameters). Both models were accessed via the official Huggingface implementation `facebook/esm1b t33 650M UR50S` and `facebook/esm2 t33 650M UR50D,` respectively.

For each sequence, residue-level representations were extracted from the final transformer layer of the respective pretrained model. Sequence-level embeddings were computed by mean pooling across all residue token embeddings, excluding special tokens such as the beginning-of-sequence and end-of-sequence markers. This aggregation yields a single contextualized representation that captures global sequence information while preserving residue-informed contextual signals. The resulting fixed-length embedding vectors have a dimensionality of 1280 for both ESM-1b and ESM-2 (650M parameter models).

Sequences exceeding the maximum token length supported by the respective model were discarded. For the PETase dataset, sequences longer than 1024 amino acids were excluded to avoid multimeric or multidomain proteins inconsistent with canonical hydrolase architectures.

Embedding extraction was performed using a NVIDIA A40 GPU with a CUDA version 12.2. Batch processing was applied with batch size 1. All embedding parameters were specified in the corresponding config.yml file to ensure reproducibility.

### 4.3 Clustering and Dimensionality Reduction

Clustering was performed directly in the high-dimensional embedding space using Hierarchical Density-Based Spatial Clustering of Applications with Noise (HDBSCAN) [McInnes et al., 2017]. Pairwise distances were computed using the default euclidean metric on the 1280-dimensional embedding vectors. HDBSCAN was selected due to its ability to identify clusters of variable density while explicitly labeling low-density points as noise, thereby avoiding the need to predefine the number of clusters.

For the LOV dataset, clustering was conducted with `min_samples`=10 and min cluster size=15. For the full PETase dataset, parameters were set to `min_samples`=10 and `min_cluster_size`=50 to account for the substantially larger dataset size. In the refined PETase subregion analysis, clustering was performed with `min_samples`=2 and min cluster size=3 to enable fine-grained neighborhood resolution. All clustering configurations were defined in the corresponding `config.yml` files to ensure reproducibility.

Nonlinear dimensionality reduction for visualization was performed using Uniform Manifold Approximation and Projection (UMAP). UMAP was applied to the same embedding vectors used for clustering, using `n_neighbors`=15 and `random_state`=42. The euclidean metric was used consistently to maintain compatibility with the clustering step. UMAP projections were used exclusively for visualization and exploratory analysis, while all quantitative clustering and connectivity reconstruction were computed in the original embedding space.

### 4.4 Connectivity and Hierarchical Reconstruction

Hierarchical relationships between sequences were reconstructed directly in the original embedding space. Pairwise distances between embedding vectors were computed using the euclidean metric, consistent with the clustering procedure. Based on this distance matrix, agglomerative hierarchical clustering was performed using the leaf linkage criterion. The resulting hierarchical tree encodes nested relationships across the full similarity continuum without requiring predefined similarity thresholds.

To recover explicit connectivity between embedding-defined neighborhoods, a Minimum Spanning Tree (MST) was derived from the hierarchical clustering structure. Specifically, the MST was constructed from the pairwise distance matrix in embedding space using Prims algorithm. This procedure yields a connected acyclic graph that preserves the minimal set of edges required to connect all sequences while minimizing total edge weight under the chosen distance metric. Importantly, both hierarchical clustering and MST reconstruction were computed in the full high-dimensional embedding space rather than on the two-dimensional UMAP projection.

Hierarchical branch lengths correspond to embedding-space distances under the selected linkage criterion and were used to quantify intra- and inter-neighborhood separation during candidate refinement. All parameters related to distance computation, linkage strategy, and connectivity reconstruction were specified in the corresponding `config.yml` file to ensure reproducibility.

### 4.5 Quantitative Evaluation

To quantify the extent to which embedding-space neighborhoods reflect functional organization in the LOV dataset, we computed k-Nearest-Neighbor (kNN) agreement for selected annotation features. For each sequence *i*, kNN agreement measures the fraction of its *k* nearest neighbors that share the same annotation label. In this work, *k* = 16 was used unless stated otherwise. Formally, for a given feature, the agreement is defined as

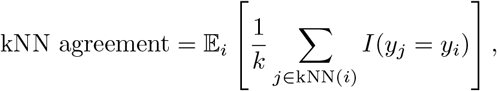

where kNN(*i*) denotes the set of the *k* nearest neighbors of sequence *i* in embedding space, *y*_*i*_ and *y*_*j*_ are the corresponding labels, and *I*(*·*) is the indicator function. The expectation is taken over all sequences in the dataset. Nearest neighbors were computed using the Minkowski distance metric in the original high-dimensional embedding space.

Because kNN agreement can be influenced by class imbalance, a permutation-based null model was constructed by randomly shuffling annotation labels while preserving class frequencies. This procedure destroys spatial structure while retaining label distribution, thereby providing a baseline estimate of agreement expected under random organization. The permutation process was repeated 200 times, and the mean and standard deviation of the resulting agreement scores were used as reference values. Observed kNN agreement values were interpreted relative to this permutation baseline to distinguish genuine embedding-space structure from effects driven purely by label imbalance.

### 4.6 Sequence and Structural Validation

To compare embedding-derived relationships with classical sequence- and structure-based similarity measures, a representative subset of seven sequences sampled from the PETase-proximal embedding-defined neighborhood was analyzed.

Multiple sequence alignment (MSA) was performed using Clustal Omega (EMBL-EBI webserver) with default parameters. Pairwise sequence identities were computed from the resulting alignment matrix. The complete alignment is provided in the Supporting Information. Sequence identity values were used to construct an all-vs-all similarity heatmap for comparison with embedding-derived relationships.

Predicted protein structures for the selected sequences were obtained from the AlphaFold Protein Structure Database (release v6). Structural superposition and pairwise structural comparisons were performed using the TM-align webserver with default parameters. Root mean square deviation (RMSD, in Å) values were extracted from the resulting pairwise alignments to generate a structural distance matrix.

Structural overlays were visualized using MOE (Molecular Operating Environment (MOE), 2024.06; Chemical Computing Group ULC, 1010 Sherbrooke St. West, Suite 910, Montreal, QC, Canada, H3A 2R7, 2024.). Superposition was performed using the default structural alignment protocol based on the BLOSUM62 substitution matrix. RMSD-based color mapping was applied to highlight regions of structural deviation, with residues exceeding 2.5 Å deviation colored in red. Non-alignable secondary structure elements of the negative control were visualized separately to emphasize structural divergence relative to embedding-defined candidates.

## Conflict of interest statement

The authors declare no conflict of interests.

### AI Statement

As non-native English speakers, the authors used ChatGPT (OpenAI, GPT) to improve the English language and stylistic tone of the manuscript. The tool was used primarily for paraphrasing and editing rather than for generating scientific content. All text was carefully reviewed and revised by the authors, who take full responsibility for the final version of the manuscript.

## Author contributions statement

F.M, D.M.O and M.D.D: conceptualization. F.M, D.M.O, A.H and M.D.D: methodology. F.M, A.K and D.M.O: validation. F.M: investigation. F.M, D.M.O, A.K, A.H and M.D.D: writing, review and editing.

M.D.D: supervision, funding resources, and project administration. All authors have read and agreed to the submission of the manuscript.

## Codes and data availability statement

The SelectZyme source code is publicly available to support local use via the GitHub repository https://github.com/ipb-halle/SelectZyme. Information about the usage and deployment are given in the repository and the Supplemental Information Section S3. The case studies presented in this paper can be explored through a public web interface https://biocloud.ipb-halle.de/selectzyme/, and the corresponding website code is available at GitHub https://github.com/ipb-halle/SelectZyme-app. Unless otherwise stated, datasets were retrieved from UniProt (release 2025 03). Configured datasets for the examples shown in the publication and on the website are available at huggingface https://huggingface.co/datasets/davari-group/selectzyme-app-data.

## Acknowledgments

The authors would like to express gratitude to Dr. Martin Dippe and Lilly Eger for frequent and constructive feedback on usability and utility in real world experimental mining scenarios which we tried to constantly integrate into the tool. For assistance on the software level, technical expertise and product deployment we like to express big gratitude to Fabian Mauz, Frank Lange and Dr. Frank Broda. A.H. acknowledges support by a Ph.D. scholarship from the Ministry of Higher Education of the Arab Republic of Egypt. Additional support from EU COST Action CA21162 (COZYME) is also acknowledged.

## Funding

This work was funded by the Deutsche Forschungsgemeinschaft (DFG, German Research Foundation) - within the Priority Program Molecular Machine Learning SPP2363 (Project Number 497207454) (M.D.D.).

## Supporting Information

The Supporting Information contains additional sections describing practical considerations for enzyme mining with SelectZyme, a brief overview of state-of-the-art enzyme mining approaches, the design rationale behind SelectZyme, further representative applications, and additional figures supporting validation of the presented workflow.

## Notes

https://github.com/ipb-halle/SelectZyme

https://github.com/ipb-halle/SelectZyme-app

https://biocloud.ipb-halle.de/selectzyme/

### Competing Interest Statement

The authors have declared no competing interest.

## References

Peter K Robinson. Enzymes: principles and biotechnological applications. Essays in biochemistry, 59:1, 2015.

Hyunbin Kim, Rachel Seongeun Kim, Milot Mirdita, and Martin Steinegger. Structural motif search across the protein-universe with folddisco. bioRxiv, pages 2025–07, July 2025. doi: 10.1101/2025.07.06.663357.

Jingi Yeo, Yewon Han, Nicola Bordin, Andy M. Lau, Shaun M. Kandathil, Hyunbin Kim, Eli Levy Karin, Milot Mirdita, David T. Jones, Christine Orengo, and Martin Steinegger. Metagenomic-scale analysis of the predicted protein structure universe. bioRxiv, pages 2025–04, April 2025. doi: 10.1101/2025.04.23.650224.

Janani Durairaj, Andrew M. Waterhouse, Toomas Mets, Tetiana Brodiazhenko, Minhal Abdullah, Gabriel Studer, Gerardo Tauriello, Mehmet Akdel, Antonina Andreeva, Alex Bateman, Tanel Tenson, Vasili Hauryliuk, Torsten Schwede, and Joana Pereira. Uncovering new families and folds in the natural protein universe. Nature, 622(7983):646–653, September 2023. ISSN 1476-4687. doi: 10.1038/s41586-023-06622-3.

Kevin K. Yang, Sarah Alamdari, Alex J. Lee, Kaeli Kaymak-Loveless, Samir Char, Garyk Brixi, Carles Domingo-Enrich, Chentong Wang, Suyue Lyu, Nicolo Fusi, Neil Tenenholtz, and Ava P. Amini. The dayhoff atlas: scaling sequence diversity for improved protein generation. bioRxiv, pages 2025–07, July 2025. doi: 10.1101/2025.07.21.665991.

Inigo Barrio-Hernandez, Jingi Yeo, Jürgen Jänes, Milot Mirdita, Cameron L. M. Gilchrist, Tanita Wein, Mihaly Varadi, Sameer Velankar, Pedro Beltrao, and Martin Steinegger. Clustering-predicted structures at the scale of the known protein universe. Nature, pages 637–645, sep 2023. doi: 10.1038/s41586-023-06510-w.

Felix Moorhoff, Yanzi Zhang, Sizhe Qiu, Wenjuan Dong, David Medina-Ortiz, Jing Zhao, and Mehdi D. Davari. Machine learning-driven enzyme mining: Opportunities, challenges, and future perspectives. ACS Catalysis, pages 12–30, December 2025. ISSN 2155-5435. doi: 10.1021/acscatal.5c04814.

Alex Bateman, Maria-Jesus Martin, Sandra Orchard, Michele Magrane, Aduragbemi Adesina, Shadab Ahmad, Emily H Bowler-Barnett, Hema Bye-A-Jee, David Carpentier, Paul Denny, Jun Fan, Penelope Garmiri, Leonardo Jose da Costa Gonzales, Abdulrahman Hussein, Alexandr Ignatchenko, Giuseppe Insana, Rizwan Ishtiaq, Vishal Joshi, Dushyanth Jyothi, Swaathi Kandasaamy, Antonia Lock, Aurelien Luciani, Jie Luo, Yvonne Lussi, Juan Sebastian Martinez Marin, Pedro Raposo, Daniel L Rice, Rafael Santos, Elena Speretta, James Stephenson, Prabhat Totoo, Nidhi Tyagi, Nadya Urakova, Preethi Vasudev, Kate Warner, Supun Wijerathne, Conny Wing-Heng Yu, Rossana Zaru, Alan J Bridge, Lucila Aimo, Ghislaine Argoud-Puy, Andrea H Auchincloss, Kristian B Axelsen, Parit Bansal, Delphine Baratin, Teresa M Batista Neto, Marie-Claude Blatter, Jerven T Bolleman, Emmanuel Boutet, Lionel Breuza, Blanca Cabrera Gil, Cristina Casals-Casas, Kamal Chikh Echioukh, Elisabeth Coudert, Beatrice Cuche, Edouard de Castro, Anne Estreicher, Maria L Famiglietti, Marc Feuermann, Elisabeth Gasteiger, Pascale Gaudet, Sebastien Gehant, Vivienne Gerritsen, Arnaud Gos, Nadine Gruaz, Chantal Hulo, Nevila Hyka-Nouspikel, Florence Jungo, Arnaud Kerhornou, Philippe Le Mercier, Damien Lieberherr, Patrick Masson, Anne Morgat, Salvo Paesano, Ivo Pedruzzi, Sandrine Pilbout, Lucille Pourcel, Sylvain Poux, Monica Pozzato, Manuela Pruess, Nicole Redaschi, Catherine Rivoire, Christian J A Sigrist, Karin Sonesson, Shyamala Sundaram, Anastasia Sveshnikova, Cathy H Wu, Cecilia N Arighi, Chuming Chen, Yongxing Chen, Hongzhan Huang, Kati Laiho, Minna Lehvaslaiho, Peter McGarvey, Darren A Natale, Karen Ross, CR Vinayaka, Yuqi Wang, and Jian Zhang. Uniprot: the universal protein knowledgebase in 2025. Nucleic Acids Research, pages 2699–2699, November 2024. ISSN 1362-4962. doi: 10.1093/nar/gkae1010.

Roshan Maroti Shinde, Sheetanshu Gupta, Dhirendra Kuma, and Mohammad Javed Ansar. Enzyme annotation and pathway reconstruction with machine learning. In Genomic Intelligence, pages 119–136. CRC Press, 2024.

Lei Zhou, Chunmeng Tao, Xiaolin Shen, Xinxiao Sun, Jia Wang, and Qipeng Yuan. Unlocking the potential of enzyme engineering via rational computational design strategies. Biotechnology Advances, page 108376, 2024.

Jason Yang, Francesca-Zhoufan Li, and Frances H. Arnold. Opportunities and challenges for machine learning-assisted enzyme engineering. ACS Central Science, 10(2):226–241, February 2024. ISSN 2374-7951. doi: 10.1021/acscentsci.3c01275.

Eric W Sayers, Jeffrey Beck, Evan E Bolton, J Rodney Brister, Jessica Chan, Ryan Connor, Michael Feldgarden, Anna M Fine, Kathryn Funk, Jinna Hoffman, Sivakumar Kannan, Christopher Kelly, William Klimke, Sunghwan Kim, Stacy Lathrop, Aron Marchler-Bauer, Terence D Murphy, Chris O’Sullivan, Erin Schmieder, Yuriy Skripchenko, Adam Stine, Francoise Thibaud-Nissen, Jiyao Wang, Jian Ye, Erin Zellers, Valerie A Schneider, and Kim D Pruitt. Database resources of the national center for biotechnology information in 2025. Nucleic Acids Research, 53(D1):D20–D29, November 2024. ISSN 1362-4962. doi: 10.1093/nar/gkae979.

Lorna Richardson, Ben Allen, Germana Baldi, Martin Beracochea, Maxwell L Bileschi, Tony Burdett, Josephine Burgin, Juan Caballero-Pérez, Guy Cochrane, Lucy J Colwell, Tom Curtis, Alejandra Escobar-Zepeda, Tatiana A Gurbich, Varsha Kale, Anton Korobeynikov, Shriya Raj, Alexander B Rogers, Ekaterina Sakharova, Santiago Sanchez, Darren J Wilkinson, and Robert D Finn. MGnify: the microbiome sequence data analysis resource in 2023. Nucleic Acids Research, 51(D1):D753–D759, dec 2022. doi: 10.1093/nar/gkac1080.

Liang Hong, Zhihang Hu, Siqi Sun, Xiangru Tang, Jiuming Wang, Qingxiong Tan, Liangzhen Zheng, Sheng Wang, Sheng Xu, Irwin King, Mark Gerstein, and Yu Li. Fast, sensitive detection of protein homologs using deep dense retrieval. Nature Biotechnology, pages 983–995, August 2024. ISSN 1546-1696. doi: 10.1038/s41587-024-02353-6.

Tymor Hamamsy, James T. Morton, Robert Blackwell, Daniel Berenberg, Nicholas Carriero, Vladimir Gligorijevic, Charlie E. M. Strauss, Julia Koehler Leman, Kyunghyun Cho, and Richard Bonneau. Protein remote homology detection and structural alignment using deep learning. Nature Biotechnology, 42(6):975–985, sep 2023. ISSN 1546-1696. doi: 10.1038/s41587-023-01917-2.

Stephen F. Altschul, Warren Gish, Webb Miller, Eugene W. Myers, and David J. Lipman. Basic local alignment search tool. Journal of Molecular Biology, 215(3):403–410, October 1990. ISSN 0022-2836. doi: 10.1016/s0022-2836(05)80360-2.

Ying Huang, Beifang Niu, Ying Gao, Limin Fu, and Weizhong Li. CD-HIT suite: a web server for clustering and comparing biological sequences. Bioinformatics, 26(5):680–682, jan 2010. doi: 10.1093/bioinformatics/btq003.

Milot Mirdita, Martin Steinegger, and Johannes Söding. Mmseqs2 desktop and local web server app for fast, interactive sequence searches. Bioinformatics, 35(16):2856–2858, January 2019. ISSN 1367-4811. doi: 10.1093/bioinformatics/bty1057.

M Ester, H Kriegel, J Sander, and Xiaowei Xu. A density-based algorithm for discovering clusters in large spatial databases with noise. KDD, pages 226–231, August 1996.

Baris E. Suzek, Hongzhan Huang, Peter McGarvey, Raja Mazumder, and Cathy H. Wu. Uniref: comprehensive and non-redundant uniprot reference clusters. Bioinformatics, 23(10):1282–1288, March 2007. ISSN 1367-4803. doi: 10.1093/bioinformatics/btm098.

K. D. Pruitt. Ncbi reference sequence (refseq): a curated non-redundant sequence database of genomes, transcripts and proteins. Nucleic Acids Research, 33(Database issue):D501–D504, December 2004. ISSN 1362-4962. doi: 10.1093/nar/gki025.

Nuala A. O’Leary, Mathew W. Wright, J. Rodney Brister, Stacy Ciufo, Diana Haddad, Rich McVeigh, Bhanu Rajput, Barbara Robbertse, Brian Smith-White, Danso Ako-Adjei, Alexander Astashyn, Azat Badretdin, Yiming Bao, Olga Blinkova, Vyacheslav Brover, Vyacheslav Chetvernin, Jinna Choi, Eric Cox, Olga Ermolaeva, Catherine M. Farrell, Tamara Goldfarb, Tripti Gupta, Daniel Haft, Eneida Hatcher, Wratko Hlavina, Vinita S. Joardar, Vamsi K. Kodali, Wenjun Li, Donna Maglott, Patrick Masterson, Kelly M. McGarvey, Michael R. Murphy, Kathleen O’Neill, Shashikant Pujar, Sanjida H. Rangwala, Daniel Rausch, Lillian D. Riddick, Conrad Schoch, Andrei Shkeda, Susan S. Storz, Hanzhen Sun, Francoise Thibaud-Nissen, Igor Tolstoy, Raymond E. Tully, Anjana R. Vatsan, Craig Wallin, David Webb, Wendy Wu, Melissa J. Landrum, Avi Kimchi, Tatiana Tatusova, Michael DiCuccio, Paul Kitts, Terence D. Murphy, and Kim D. Pruitt. Reference sequence (RefSeq) database at NCBI: current status, taxonomic expansion, and functional annotation. Nucleic Acids Research, 44(D1):D733–D745, nov 2015. doi: 10.1093/nar/gkv1189.

Qingyu Chen, Yu Wan, Yang Lei, Justin Zobel, and Karin Verspoor. Evaluation of cd-hit for constructing non-redundant databases. In 2016 IEEE International Conference on Bioinformatics and Biomedicine (BIBM), pages 703–706. IEEE, December 2016. doi: 10.1109/bibm.2016.7822604.

Rémi Zallot, Nils Oberg, and John A. Gerlt. The EFI web resource for genomic enzymology tools: Leveraging protein, genome, and metagenome databases to discover novel enzymes and metabolic pathways. Biochemistry, 58(41):4169–4182, sep 2019. doi: 10.1021/acs.biochem.9b00735.

Janine N. Copp, Dave W. Anderson, Eyal Akiva, Patricia C. Babbitt, and Nobuhiko Tokuriki. Exploring the sequence, function, and evolutionary space of protein superfamilies using sequence similarity networks and phylogenetic reconstructions. In Methods in Enzymology, pages 315–347. Elsevier, 2019. doi: 10.1016/bs.mie.2019.03.015.

Bastian Volker Helmut Hornung and Nicolas Terrapon. An objective criterion to evaluate sequence-similarity networks helps in dividing the protein family sequence space. PLOS Computational Biology, 19(8):e1010881, 2023. doi: 10.1101/2022.04.19.488343.

John Z Chen, Barnabas Gall, Sacha B Pulsford, Nobuhiko Tokuriki, and Colin J Jackson. Exploring large protein sequence space through homology- and representation-based hierarchical clustering. Molecular Biology and Evolution, 42(6):msaf136, June 2025. ISSN 1537-1719. doi: 10.1093/molbev/msaf136.

John A. Gerlt, Jason T. Bouvier, Daniel B. Davidson, Heidi J. Imker, Boris Sadkhin, David R. Slater, and Katie L. Whalen. Enzyme function initiative-enzyme similarity tool (efi-est): A web tool for generating protein sequence similarity networks. Biochimica et Biophysica Acta (BBA) - Proteins and Proteomics, 1854(8):1019–1037, August 2015. ISSN 1570-9639. doi: 10.1016/j.bbapap.2015.04.015.

Jiri Hon, Simeon Borko, Jan Stourac, Zbynek Prokop, Jaroslav Zendulka, David Bednar, Tomas Martinek, and Jiri Damborsky. EnzymeMiner: automated mining of soluble enzymes with diverse structures, catalytic properties and stabilities. Nucleic Acids Research, 48(W1):W104–W109, may 2020. doi: 10.1093/nar/gkaa372.

Alexander Rives, Joshua Meier, Tom Sercu, Siddharth Goyal, Zeming Lin, Jason Liu, Demi Guo, Myle Ott, C. Lawrence Zitnick, Jerry Ma, and Rob Fergus. Biological structure and function emerge from scaling unsupervised learning to 250 million protein sequences. Proceedings of the National Academy of Sciences, 118(15):e2016239118, apr 2021. doi: 10.1073/pnas.2016239118.

Kyra Erckert and Burkhard Rost. Assessing the role of evolutionary information for enhancing protein language model embeddings. Scientific Reports, 14(1):20692, September 2024. ISSN 2045-2322. doi: 10.1038/s41598-024-71783-8.

Zeming Lin, Halil Akin, Roshan Rao, Brian Hie, Zhongkai Zhu, Wenting Lu, Nikita Smetanin, Robert Verkuil, Ori Kabeli, Yaniv Shmueli, Allan dos Santos Costa, Maryam Fazel-Zarandi, Tom Sercu, Salvatore Candido, and Alexander Rives. Evolutionary-scale prediction of atomic-level protein structure with a language model. Science, 379(6637):1123–1130, March 2023. ISSN 1095-9203. doi: 10.1126/science.ade2574.

Michael Heinzinger, Konstantin Weissenow, Joaquin Gomez Sanchez, Adrian Henkel, Milot Mirdita, Martin Steinegger, and Burkhard Rost. Bilingual language model for protein sequence and structure. NAR Genomics and Bioinformatics, 6(4):lqae150, September 2024. ISSN 2631-9268. doi: 10.1093/nargab/lqae150.

Tobias Senoner, Ivan Koludarov, Joshua Günther, Amarda Shehu, Burkhard Rost, and Yana Bromberg. Which plm to choose? bioRxiv, pages 2025–10, October 2025a. doi: 10.1101/2025.10.30.685515.

Jin Su, Chenchen Han, Yuyang Zhou, Junjie Shan, Xibin Zhou, and Fajie Yuan. Saprot: Protein language modeling with structure-aware vocabulary. bioRxiv, pages 2023–10, October 2023. doi: 10.1101/2023.10.01.560349.

Ahmed Elnaggar, Michael Heinzinger, Christian Dallago, Ghalia Rehawi, Yu Wang, Llion Jones, Tom Gibbs, Tamas Feher, Christoph Angerer, Martin Steinegger, Debsindhu Bhowmik, and Burkhard Rost. Prottrans: Toward understanding the language of life through self-supervised learning. IEEE Transactions on Pattern Analysis and Machine Intelligence, 44(10):7112–7127, October 2022. ISSN 1939-3539. doi: 10.1109/tpami.2021.3095381.

Tobias Senoner, Tobias Olenyi, Michael Heinzinger, Anton Spannagl, George Bouras, Burkhard Rost, and Ivan Koludarov. Protspace: A tool for visualizing protein space. Journal of Molecular Biology, 437(15):168940, August 2025b. ISSN 0022-2836. doi: 10.1016/j.jmb.2025.168940.

Serina L. Robinson, Jörn Piel, and Shinichi Sunagawa. A roadmap for metagenomic enzyme discovery. Natural Product Reports, 38(11):1994–2023, 2021. doi: 10.1039/d1np00006c.

Elizabeth Fahsbender, Alma Andersson, Jeremy Ash, Polina Binder, Daniel Burkhardt, Benjamin Chang, Georg K. Gerber, Anthony Gitter, Patrick Godau, Ankit Gupta, Genevieve Haliburton, Siyu He, Trey Ideker, Ivana Jelic, Aly Khan, Yang-Joon Kim, Aditi Krishnapriyan, Jon M. Laurent, Tianyu Liu, Emma Lundberg, Shalin B. Mehta, Rob Moccia, Angela Oliveira Pisco, Katherine S. Pollard, Suresh Ramani, Julio Saez-Rodriguez, Yasin Senbabaoglu, Elana Simon, Srinivasan Sivanandan, Gustavo Stolovitzky, Marc Valer, Bo Wang, Xikun Zhang, James Zou, and Katrina Kalantar. Benchmarking and evaluation of ai models in biology: Outcomes and recommendations from the czi virtual cells workshop, 2025.

Judith Bernett, David B. Blumenthal, Dominik G. Grimm, Florian Haselbeck, Roman Joeres, Olga V. Kalinina, and Markus List. Guiding questions to avoid data leakage in biological machine learning applications. Nature Methods, 21(8):1444–1453, August 2024. ISSN 1548-7105. doi: 10.1038/s41592-024-02362-y.

Anton Bushuiev, Roman Bushuiev, Jiri Sedlar, Tomas Pluskal, Jiri Damborsky, Stanislav Mazurenko, and Josef Sivic. Revealing data leakage in protein interaction benchmarks, 2024.

Sayash Kapoor and Arvind Narayanan. Leakage and the reproducibility crisis in machine-learning-based science. Patterns, 4(9):100804, September 2023. ISSN 2666-3899. doi: 10.1016/j.patter.2023.100804.

Anna Klimovskaia Susmelj, Yani Ren, Yann Vander Meersche, Jean-Christophe Gelly, and Tatiana Galochkina. Poincaré maps for visualization of large protein families. Briefings in Bioinformatics, 24(3):bbad103, March 2023. ISSN 1477-4054. doi: 10.1093/bib/bbad103.

J. C. Gower and G. J. S. Ross. Minimum spanning trees and single linkage cluster analysis. Applied Statistics, 18(1):54, 1969. ISSN 0035-9254. doi: 10.2307/2346439.

Kamalika Chaudhuri, Sanjoy Dasgupta, Samory Kpotufe, and Ulrike von Luxburg. Consistent procedures for cluster tree estimation and pruning. IEEE Transactions on Information Theory, 60(12):7900–7912, December 2014. ISSN 1557-9654. doi: 10.1109/tit.2014.2361055.

Daniel Probst and Jean-Louis Reymond. Visualization of very large high-dimensional data sets as minimum spanning trees. Journal of Cheminformatics, 12(1):12, February 2020. ISSN 1758-2946. doi: 10.1186/s13321-020-0416-x.

Kristof Theys, Philippe Lemey, Anne-Mieke Vandamme, and Guy Baele. Advances in visualization tools for phylogenomic and phylodynamic studies of viral diseases. Frontiers in Public Health, 7, August 2019. ISSN 2296-2565. doi: 10.3389/fpubh.2019.00208.

Spencer T. Glantz, Eric J. Carpenter, Michael Melkonian, Kevin H. Gardner, Edward S. Boyden, Gane Ka-Shu Wong, and Brian Y. Chow. Functional and topological diversity of lov domain photoreceptors. Proceedings of the National Academy of Sciences, 113(11):E1442–E1451, February 2016. ISSN 1091-6490. doi: 10.1073/pnas.1509428113.

Benita Kopka, Kathrin Magerl, Anton Savitsky, Mehdi D. Davari, Katrin Röllen, Marco Bocola, Bernhard Dick, Ulrich Schwaneberg, Karl-Erich Jaeger, and Ulrich Krauss. Electron transfer pathways in a light, oxygen, voltage (lov) protein devoid of the photoactive cysteine. Scientific Reports, 7(1):13346, October 2017. ISSN 2045-2322. doi: 10.1038/s41598-017-13420-1.

Ren Wei, Peter Westh, Gert Weber, Lars M. Blank, and Uwe T. Bornscheuer. Standardization guidelines and future trends for pet hydrolase research. Nature Communications, 16(1):4684, May 2025a. ISSN 2041-1723. doi: 10.1038/s41467-025-60016-9.

Patrick C. F. Buchholz, Golo Feuerriegel, Hongli Zhang, Pablo Perez-Garcia, Lena-Luisa Nover, Jennifer Chow, Wolfgang R. Streit, and Jürgen Pleiss. Plastics degradation by hydrolytic enzymes: The plastics-active enzymes database— pazy. Proteins: Structure, Function, and Bioinformatics, 90(7):1443–1456, February 2022. ISSN 1097-0134. doi: 10.1002/prot.26325.

Hogyun Seo, Hwaseok Hong, Jiyoung Park, Seul Hoo Lee, Dongwoo Ki, Aejin Ryu, Hye-Young Sagong, and Kyung-Jin Kim. Landscape profiling of pet depolymerases using a natural sequence cluster framework. Science, 387(6729):eadp5637, January 2025. ISSN 1095-9203. doi: 10.1126/science.adp5637.

Onur Turak, Andreas Gagsteiger, Ashank Upadhyay, Mark Kriegel, Peter Salein, Stefanie Böhnke-Brandt, Seema Agarwal, Erik Borchert, and Birte Höcker. A third type of petase from the marine halopseudomonas lineage. Protein Science, 34(10), September 2025. ISSN 1469-896X. doi: 10.1002/pro.70305.

David Medina-Ortiz, Diego Alvarez-Saravia, Nicole Soto-García, Diego Sandoval-Vargas, Jacqueline Aldridge, Sebastián Rodríguez, Barbara Andrews, Juan A. Asenjo, and Anamaría Daza. Protein language models accelerate the discovery of plastic-degrading enzymes. bioRxiv, pages 2025–02, February 2025. doi: 10.1101/2025.02.09.637306.

Ren Wei, Gert Weber, Lars M. Blank, and Uwe T. Bornscheuer. Process insights for harnessing biotechnology for plastic depolymerization. Nature Chemical Engineering, 2(2):110–117, February 2025b. ISSN 2948-1198. doi: 10.1038/s44286-024-00171-w.

Burkhard Rost. Twilight zone of protein sequence alignments. Protein Engineering, Design and Selection, 12(2):85–94, February 1999. ISSN 1741-0126. doi: 10.1093/protein/12.2.85.

Frances Ding and Jacob Steinhardt. Protein language models are biased by unequal sequence sampling across the tree of life. March 2024. doi: 10.1101/2024.03.07.584001.

Andreas Bjerregaard, Peter Mørch Groth, Søren Hauberg, Anders Krogh, and Wouter Boomsma. Foundation models of protein sequences: A brief overview. Current Opinion in Structural Biology, 91:103004, April 2025. ISSN 0959-440X. doi: 10.1016/j.sbi.2025.103004.

Klemens Flöge, Srisruthi Udayakumar, Johanna Sommer, Marie Piraud, Stefan Kesselheim, Vincent Fortuin, Stephan Günnemann, Karel J. van der Weg, Holger Gohlke, Erinc Merdivan, and Alina Bazarova. Oneprot: Towards multi-modal protein foundation models via latent space alignment of sequence, structure, binding sites and text encoders. PLOS Computational Biology, 21(11):e1013679, November 2025. ISSN 1553-7358. doi: 10.1371/journal.pcbi.1013679.

Jin Su, Yan He, Shiyang You, Shiyu Jiang, Xibin Zhou, Xuting Zhang, Yuxuan Wang, Xining Su, Igor Tolstoy, Xing Chang, Hongyuan Lu, and Fajie Yuan. A trimodal protein language model enables advanced protein searches. Nature Biotechnology, October 2025. ISSN 1546-1696. doi: 10.1038/s41587-025-02836-0.

Dehua Peng, Zhipeng Gui, Wenzhang Wei, Fa Li, Jie Gui, Huayi Wu, and Jianya Gong. Sampling-enabled scalable manifold learning unveils the discriminative cluster structure of high-dimensional data. Nature Machine Intelligence, September 2025. ISSN 2522-5839. doi: 10.1038/s42256-025-01112-9.

Victor Gambarini, Olga Pantos, Joanne M Kingsbury, Louise Weaver, Kim M Handley, and Gavin Lear. Plasticdb: a database of microorganisms and proteins linked to plastic biodegradation. Database, 2022:baac008, January 2022. ISSN 1758-0463. doi: 10.1093/database/baac008.

Leland McInnes, John Healy, and Steve Astels. hdbscan: Hierarchical density based clustering. The Journal of Open Source Software, 2(11):205, March 2017. ISSN 2475-9066. doi: 10.21105/joss.00205.

## References

Altschul, S. F., Gish, W., Miller, W., Myers, E. W., and Lipman, D. J. (1990). Basic local alignment search tool. Journal of Molecular Biology, 215(3):403–410.

Barrio-Hernandez, I., Yeo, J., Jänes, J., Mirdita, M., Gilchrist, C. L. M., Wein, T., Varadi, M., Velankar, S., Beltrao, P., and Steinegger, M. (2023). Clustering-predicted structures at the scale of the known protein universe. Nature, pages 637–645.

Bateman, A., Martin, M.-J., Orchard, S., Magrane, M., Adesina, A., Ahmad, S., Bowler-Barnett, E. H., Bye-A-Jee, H., Carpentier, D., Denny, P., Fan, J., Garmiri, P., Gonzales, L. J. d. C., Hussein, A., Ignatchenko, A., Insana, G., Ishtiaq, R., Joshi, V., Jyothi, D., Kandasaamy, S., Lock, A., Luciani, A., Luo, J., Lussi, Y., Marin, J. S. M., Raposo, P., Rice, D. L., Santos, R., Speretta, E., Stephenson, J., Totoo, P., Tyagi, N., Urakova, N., Vasudev, P., Warner, K., Wijerathne, S., Yu, C. W.-H., Zaru, R., Bridge, A. J., Aimo, L., Argoud-Puy, G., Auchincloss, A. H., Axelsen, K. B., Bansal, P., Baratin, D., Batista Neto, T. M., Blatter, M.-C., Bolleman, J. T., Boutet, E., Breuza, L., Gil, B. C., Casals-Casas, C., Echioukh, K. C., Coudert, E., Cuche, B., de Castro, E., Estreicher, A., Famiglietti, M. L., Feuermann, M., Gasteiger, E., Gaudet, P., Gehant, S., Gerritsen, V., Gos, A., Gruaz, N., Hulo, C., Hyka-Nouspikel, N., Jungo, F., Kerhornou, A., Mercier, P. L., Lieberherr, D., Masson, P., Morgat, A., Paesano, S., Pedruzzi, I., Pilbout, S., Pourcel, L., Poux, S., Pozzato, M., Pruess, M., Redaschi, N., Rivoire, C., Sigrist, C. J. A., Sonesson, K., Sundaram, S., Sveshnikova, A., Wu, C. H., Arighi, C. N., Chen, C., Chen, Y., Huang, H., Laiho, K., Lehvaslaiho, M., McGarvey, P., Natale, D. A., Ross, K., Vinayaka, C. R., Wang, Y., and Zhang, J. (2024). Uniprot: the universal protein knowledgebase in 2025. Nucleic Acids Research, pages 2699–2699.

Bernett, J., Blumenthal, D. B., Grimm, D. G., Haselbeck, F., Joeres, R., Kalinina, O. V., and List, M. (2024). Guiding questions to avoid data leakage in biological machine learning applications. Nature Methods, 21(8):1444–1453.

Bjerregaard, A., Groth, P. M., Hauberg, S., Krogh, A., and Boomsma, W. (2025). Foundation models of protein sequences: A brief overview. Current Opinion in Structural Biology, 91:103004.

Buchholz, P. C. F., Feuerriegel, G., Zhang, H., Perez-Garcia, P., Nover, L., Chow, J., Streit, W. R., and Pleiss, J. (2022). Plastics degradation by hydrolytic enzymes: The plastics-active enzymes database— pazy. Proteins: Structure, Function, and Bioinformatics, 90(7):1443–1456.

Bushuiev, A., Bushuiev, R., Sedlar, J., Pluskal, T., Damborsky, J., Mazurenko, S., and Sivic, J. (2024). Revealing data leakage in protein interaction benchmarks.

Chaudhuri, K., Dasgupta, S., Kpotufe, S., and von Luxburg, U. (2014). Consistent procedures for cluster tree estimation and pruning. IEEE Transactions on Information Theory, 60(12):7900–7912.

Chen, J. Z., Gall, B., Pulsford, S. B., Tokuriki, N., and Jackson, C. J. (2025). Exploring large protein sequence space through homology- and representation-based hierarchical clustering. Molecular Biology and Evolution, 42(6):msaf136.

Chen, Q., Wan, Y., Lei, Y., Zobel, J., and Verspoor, K. (2016). Evaluation of cd-hit for constructing non-redundant databases. In 2016 IEEE International Conference on Bioinformatics and Biomedicine (BIBM), pages 703–706. IEEE.

Copp, J. N., Anderson, D. W., Akiva, E., Babbitt, P. C., and Tokuriki, N. (2019). Exploring the sequence, function, and evolutionary space of protein superfamilies using sequence similarity networks and phylogenetic reconstructions. In Methods in Enzymology, pages 315–347. Elsevier.

Ding, F. and Steinhardt, J. (2024). Protein language models are biased by unequal sequence sampling across the tree of life.

Durairaj, J., Waterhouse, A. M., Mets, T., Brodiazhenko, T., Abdullah, M., Studer, G., Tauriello, G., Akdel, M., Andreeva, A., Bateman, A., Tenson, T., Hauryliuk, V., Schwede, T., and Pereira, J. (2023). Uncovering new families and folds in the natural protein universe. Nature, 622(7983):646–653.

Elnaggar, A., Heinzinger, M., Dallago, C., Rehawi, G., Wang, Y., Jones, L., Gibbs, T., Feher, T., Angerer, C., Steinegger, M., Bhowmik, D., and Rost, B. (2022). Prottrans: Toward understanding the language of life through self-supervised learning. IEEE Transactions on Pattern Analysis and Machine Intelligence, 44(10):7112–7127.

Erckert, K. and Rost, B. (2024). Assessing the role of evolutionary information for enhancing protein language model embeddings. Scientific Reports, 14(1):20692.

Ester, M., Kriegel, H., Sander, J., and Xu, X. (1996). A density-based algorithm for discovering clusters in large spatial databases with noise. KDD, pages 226–231.

Fahsbender, E., Andersson, A., Ash, J., Binder, P., Burkhardt, D., Chang, B., Gerber, G. K., Gitter, A., Godau, P., Gupta, A., Haliburton, G., He, S., Ideker, T., Jelic, I., Khan, A., Kim, Y.-J., Krishnapriyan, A., Laurent, J. M., Liu, T., Lundberg, E., Mehta, S. B., Moccia, R., Pisco, A. O., Pollard, K. S., Ramani, S., Saez-Rodriguez, J., Senbabaoglu, Y., Simon, E., Sivanandan, S., Stolovitzky, G., Valer, M., Wang, B., Zhang, X., Zou, J., and Kalantar, K. (2025). Benchmarking and evaluation of ai models in biology: Outcomes and recommendations from the czi virtual cells workshop.

Flöge, K., Udayakumar, S., Sommer, J., Piraud, M., Kesselheim, S., Fortuin, V., Günnemann, S., van der Weg, K. J., Gohlke, H., Merdivan, E., and Bazarova, A. (2025). Oneprot: Towards multi-modal protein foundation models via latent space alignment of sequence, structure, binding sites and text encoders. PLOS Computational Biology, 21(11):e1013679.

Gambarini, V., Pantos, O., Kingsbury, J. M., Weaver, L., Handley, K. M., and Lear, G. (2022). Plasticdb: a database of microorganisms and proteins linked to plastic biodegradation. Database, 2022:baac008.

Gerlt, J. A., Bouvier, J. T., Davidson, D. B., Imker, H. J., Sadkhin, B., Slater, D. R., and Whalen, K. L. (2015). Enzyme function initiative-enzyme similarity tool (efi-est): A web tool for generating protein sequence similarity networks. Biochimica et Biophysica Acta (BBA) - Proteins and Proteomics, 1854(8):1019–1037.

Glantz, S. T., Carpenter, E. J., Melkonian, M., Gardner, K. H., Boyden, E. S., Wong, G. K.-S., and Chow, B. Y. (2016). Functional and topological diversity of lov domain photoreceptors. Proceedings of the National Academy of Sciences, 113(11):E1442–E1451.

Gower, J. C. and Ross, G. J. S. (1969). Minimum spanning trees and single linkage cluster analysis. Applied Statistics, 18(1):54.

Hamamsy, T., Morton, J. T., Blackwell, R., Berenberg, D., Carriero, N., Gligorijevic, V., Strauss, C. E. M., Leman, J. K., Cho, K., and Bonneau, R. (2023). Protein remote homology detection and structural alignment using deep learning. Nature Biotechnology, 42(6):975–985.

Heinzinger, M., Weissenow, K., Sanchez, J. G., Henkel, A., Mirdita, M., Steinegger, M., and Rost, B. (2024). Bilingual language model for protein sequence and structure. NAR Genomics and Bioinformatics, 6(4):qae150.

Hon, J., Borko, S., Stourac, J., Prokop, Z., Zendulka, J., Bednar, D., Martinek, T., and Damborsky, J. (2020). EnzymeMiner: automated mining of soluble enzymes with diverse structures, catalytic properties and stabilities. Nucleic Acids Research, 48(W1):W104–W109.

Hong, L., Hu, Z., Sun, S., Tang, X., Wang, J., Tan, Q., Zheng, L., Wang, S., Xu, S., King, I., Gerstein, M., and Li, Y. (2024). Fast, sensitive detection of protein homologs using deep dense retrieval. Nature Biotechnology, pages 983–995.

Hornung, B. V. H. and Terrapon, N. (2023). An objective criterion to evaluate sequence-similarity networks helps in dividing the protein family sequence space. PLOS Computational Biology, 19(8):e1010881.

Huang, Y., Niu, B., Gao, Y., Fu, L., and Li, W. (2010). CD-HIT suite: a web server for clustering and comparing biological sequences. Bioinformatics, 26(5):680–682.

Kapoor, S. and Narayanan, A. (2023). Leakage and the reproducibility crisis in machine-learning-based science. Patterns, 4(9):100804.

Kim, H., Kim, R. S., Mirdita, M., and Steinegger, M. (2025). Structural motif search across the proteinuniverse with folddisco. bioRxiv, pages 2025–07.

Kopka, B., Magerl, K., Savitsky, A., Davari, M. D., Röllen, K., Bocola, M., Dick, B., Schwaneberg, U., Jaeger, K.-E., and Krauss, U. (2017). Electron transfer pathways in a light, oxygen, voltage (lov) protein devoid of the photoactive cysteine. Scientific Reports, 7(1):13346.

Lin, Z., Akin, H., Rao, R., Hie, B., Zhu, Z., Lu, W., Smetanin, N., Verkuil, R., Kabeli, O., Shmueli, Y., dos Santos Costa, A., Fazel-Zarandi, M., Sercu, T., Candido, S., and Rives, A. (2023). Evolutionary-scale prediction of atomic-level protein structure with a language model. Science, 379(6637):1123–1130.

McInnes, L., Healy, J., and Astels, S. (2017). hdbscan: Hierarchical density based clustering. The Journal of Open Source Software, 2(11):205.

Medina-Ortiz, D., Alvarez-Saravia, D., Soto-García, N., Sandoval-Vargas, D., Aldridge, J., Rodríguez, S., Andrews, B., Asenjo, J. A., and Daza, A. (2025). Protein language models accelerate the discovery of plastic-degrading enzymes. bioRxiv, pages 2025–02.

Mirdita, M., Steinegger, M., and Söding, J. (2019). Mmseqs2 desktop and local web server app for fast, interactive sequence searches. Bioinformatics, 35(16):2856–2858.

Moorhoff, F., Zhang, Y., Qiu, S., Dong, W., Medina-Ortiz, D., Zhao, J., and Davari, M. D. (2025). Machine learning-driven enzyme mining: Opportunities, challenges, and future perspectives. ACS Catalysis, pages 12–30.

O’Leary, N. A., Wright, M. W., Brister, J. R., Ciufo, S., Haddad, D., McVeigh, R., Rajput, B., Robbertse, B., Smith-White, B., Ako-Adjei, D., Astashyn, A., Badretdin, A., Bao, Y., Blinkova, O., Brover, V., Chetvernin, V., Choi, J., Cox, E., Ermolaeva, O., Farrell, C. M., Goldfarb, T., Gupta, T., Haft, D., Hatcher, E., Hlavina, W., Joardar, V. S., Kodali, V. K., Li, W., Maglott, D., Masterson, P., McGarvey, K. M., Murphy, M. R., O’Neill, K., Pujar, S., Rangwala, S. H., Rausch, D., Riddick, L. D., Schoch, C., Shkeda, A., Storz, S. S., Sun, H., Thibaud-Nissen, F., Tolstoy, I., Tully, R. E., Vatsan, A. R., Wallin, C., Webb, D., Wu, W., Landrum, M. J., Kimchi, A., Tatusova, T., DiCuccio, M., Kitts, P., Murphy, T. D., and Pruitt, K. D. (2015). Reference sequence (RefSeq) database at NCBI: current status, taxonomic expansion, and functional annotation. Nucleic Acids Research, 44(D1):D733–D745.

Peng, D., Gui, Z., Wei, W., Li, F., Gui, J., Wu, H., and Gong, J. (2025). Sampling-enabled scalable manifold learning unveils the discriminative cluster structure of high-dimensional data. Nature Machine Intelligence.

Probst, D. and Reymond, J.-L. (2020). Visualization of very large high-dimensional data sets as minimum spanning trees. Journal of Cheminformatics, 12(1):12.

Pruitt, K. D. (2004). Ncbi reference sequence (refseq): a curated non-redundant sequence database of genomes, transcripts and proteins. Nucleic Acids Research, 33(Database issue):D501–D504.

Richardson, L., Allen, B., Baldi, G., Beracochea, M., Bileschi, M. L., Burdett, T., Burgin, J., Caballero-Pérez, J., Cochrane, G., Colwell, L. J., Curtis, T., Escobar-Zepeda, A., Gurbich, T. A., Kale, V., Korobeynikov, A., Raj, S., Rogers, A. B., Sakharova, E., Sanchez, S., Wilkinson, D. J., and Finn, R. D. (2022). MGnify: the microbiome sequence data analysis resource in 2023. Nucleic Acids Research, 51(D1):D753–D759.

Rives, A., Meier, J., Sercu, T., Goyal, S., Lin, Z., Liu, J., Guo, D., Ott, M., Zitnick, C. L., Ma, J., and Fergus, R. (2021). Biological structure and function emerge from scaling unsupervised learning to 250 million protein sequences. Proceedings of the National Academy of Sciences, 118(15):e2016239118.

Robinson, P. K. (2015). Enzymes: principles and biotechnological applications. Essays in biochemistry, 59:1.

Robinson, S. L., Piel, J., and Sunagawa, S. (2021). A roadmap for metagenomic enzyme discovery. Natural Product Reports, 38(11):1994–2023.

Rost, B. (1999). Twilight zone of protein sequence alignments. Protein Engineering, Design and Selection, 12(2):85–94.

Sayers, E., Beck, J., Bolton, E., Brister, J., Chan, J., Connor, R., Feldgarden, M., Fine, A., Funk, K., Hoffman, J., Kannan, S., Kelly, C., Klimke, W., Kim, S., Lathrop, S., Marchler-Bauer, A., Murphy, T., O’Sullivan, C., Schmieder, E., Skripchenko, Y., Stine, A., Thibaud-Nissen, F., Wang, J., Ye, J., Zellers, E., Schneider, V., and Pruitt, K. (2024). Database resources of the national center for biotechnology information in 2025. Nucleic Acids Research, 53(D1):D20–D29.

Senoner, T., Koludarov, I., Günther, J., Shehu, A., Rost, B., and Bromberg, Y. (2025a). Which plm to choose? bioRxiv, pages 2025–10.

Senoner, T., Olenyi, T., Heinzinger, M., Spannagl, A., Bouras, G., Rost, B., and Koludarov, I. (2025b). Protspace: A tool for visualizing protein space. Journal of Molecular Biology, 437(15):168940.

Seo, H., Hong, H., Park, J., Lee, S. H., Ki, D., Ryu, A., Sagong, H.-Y., and Kim, K.-J. (2025). Landscape profiling of pet depolymerases using a natural sequence cluster framework. Science, 387(6729):eadp5637.

Shinde, R. M., Gupta, S., Kuma, D., and Ansar, M. J. (2024). Enzyme annotation and pathway reconstruction with machine learning. In Genomic Intelligence, pages 119–136. CRC Press.

Su, J., Han, C., Zhou, Y., Shan, J., Zhou, X., and Yuan, F. (2023). Saprot: Protein language modeling with structure-aware vocabulary. bioRxiv, pages 2023–10.

Su, J., He, Y., You, S., Jiang, S., Zhou, X., Zhang, X., Wang, Y., Su, X., Tolstoy, I., Chang, X., Lu, H., and Yuan, F. (2025). A trimodal protein language model enables advanced protein searches. Nature Biotechnology.

Susmelj, A. K., Ren, Y., Vander Meersche, Y., Gelly, J.-C., and Galochkina, T. (2023). Poincaré maps for visualization of large protein families. Briefings in Bioinformatics, 24(3):bbad103.

Suzek, B. E., Huang, H., McGarvey, P., Mazumder, R., and Wu, C. H. (2007). Uniref: comprehensive and non-redundant uniprot reference clusters. Bioinformatics, 23(10):1282–1288.

Theys, K., Lemey, P., Vandamme, A.-M., and Baele, G. (2019). Advances in visualization tools for phylogenomic and phylodynamic studies of viral diseases. Frontiers in Public Health, 7.

Turak, O., Gagsteiger, A., Upadhyay, A., Kriegel, M., Salein, P., Böhnke-Brandt, S., Agarwal, S., Borchert, E., and Höcker, B. (2025). A third type of petase from the marine halopseudomonas lineage. Protein Science, 34(10).

Wei, R., Weber, G., Blank, L. M., and Bornscheuer, U.T. (2025a). Process insights for harnessing biotechnology for plastic depolymerization. Nature Chemical Engineering, 2(2):110–117.

Wei, R., Westh, P., Weber, G., Blank, L. M., and Bornscheuer, U. T. (2025b). Standardization guidelines and future trends for pet hydrolase research. Nature Communications, 16(1):4684.

Yang, J., Li, F.-Z., and Arnold, F. H. (2024). Opportunities and challenges for machine learning-assisted enzyme engineering. ACS Central Science, 10(2):226–241.

Yang, K. K., Alamdari, S., Lee, A. J., Kaymak-Loveless, K., Char, S., Brixi, G., Domingo-Enrich, C., Wang, C., Lyu, S., Fusi, N., Tenenholtz, N., and Amini, A. P. (2025). The dayhoff atlas: scaling sequence diversity for improved protein generation. bioRxiv, pages 2025–07.

Yeo, J., Han, Y., Bordin, N., Lau, A. M., Kandathil, S. M., Kim, H., Karin, E. L., Mirdita, M., Jones, D. T., Orengo, C., and Steinegger, M. (2025). Metagenomic-scale analysis of the predicted protein structure universe. bioRxiv, pages 2025–04.

Zallot, R., Oberg, N., and Gerlt, J. A. (2019). The EFI web resource for genomic enzymology tools: Leveraging protein, genome, and metagenome databases to discover novel enzymes and metabolic pathways. Biochemistry, 58(41):4169–4182.

Zhou, L., Tao, C., Shen, X., Sun, X., Wang, J., and Yuan, Q. (2024). Unlocking the potential of enzyme engineering via rational computational design strategies. Biotechnology Advances, page 108376.

