## Supporting Information for "“Visualize, Explore, and Select”: A Protein Language Model-based Approach Enabling Navigation of Protein Sequence Space for Enzyme Discovery and Mining"

### List of Tables

**S1** List of pre-trained models available in SelectZyme . . . . . . . . . . . . . . . . 11

### Enzyme mining considerations

To provide a practical roadmap, we structure enzyme mining into the four fundamental steps: (i) *enzyme pool creation* (defining the search space via queries, signatures, or structural criteria), (ii) *sequence comparison* (quantifying relatedness across the pool), (iii) *visualization* (interpreting global structure and local neighborhoods to identify informative sub-regions), and (iv) *candidate selection* (prioritizing representative and actionable sequences for experimental follow-up) (Figure **S1**). Each step implicitly encodes constraints and objectives—such as the desired breadth of novelty, the tolerance for low-homology relationships, and practical requirements linked to organism choice, substrate scope, or process conditions—that are rarely captured by a single metric or one-size-fits-all parameter setting. For each step, we highlight common *considerations and objectives* that determine how narrowly or broadly to explore the sequence space. We also provide further *practical advice for the usage with SelectZyme*. The structure is intended to help readers conduct mining workflows that remain comprehensible and experimentally actionable, while retaining sufficient flexibility to iterate as new hypotheses, annotations, or constraints emerge during exploration.

#### Enzyme pool creation

Since enzyme pool creation can be very variable to the objective, it is not part of our proposed workflow. Nevertheless, we like to give some considerations and advice of important aspects. To form an enzyme pool, not only individual sequences of interest must be selected, but more often a strategy is performed by searching for specific functionalities or broad protein motives or folds e.g.. The characteristics of such an enzyme pool might be as versatile as its desired applications. Sequential diversity (sequence homology) plays a central role in defining how locally vs. globally a functional landscape should be explored. Additionally, depending on the target of study there can also be too many sequences included for a specific study. All in all, the pool has to be defined and selected accordingly to answer the scientific

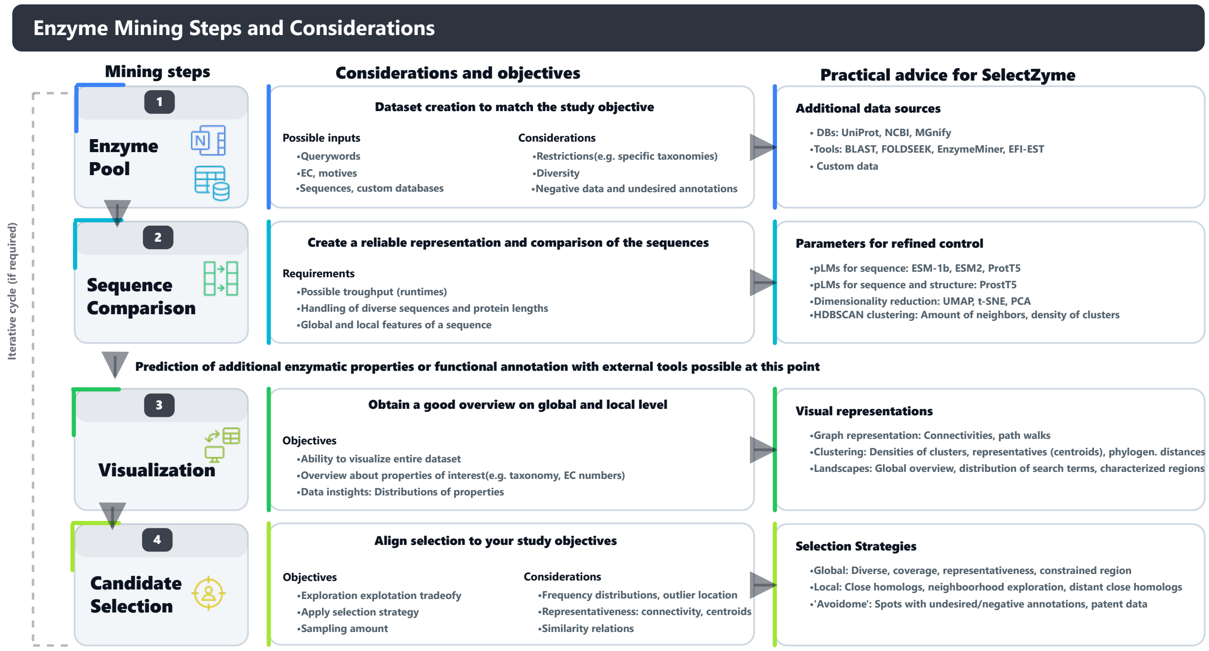

**Figure S1: Enzyme mining steps and considerations.**
Enzyme mining consists of the four main 'steps': enzyme pool creation, sequence comparison, visualization and candidate selection. Each step implicitly defines requirements and objectives that are stated in 'Considerations and objectives'. 'Practical advice' detail available computational concepts, tools or strategies to optimize enzyme mining in SelectZyme.

research questions and respect diversity, size and characteristics of the selected sequences.

Most common selection practices follow the central dogma that sequence similarity 'exhibits' similar structural folds which exhibit similar functionalities. However, if the pool needs to be defined wider, including more functionalities, or if entire superfamilies are subject of study, sequence motives, domains or entire folds can be incorporated. A narrow search (small diversity and low amount of sequences) via simple alignment-based strategies like a BLAST[2] query with default parameters can already yield the desired outcome. Next to changing alignment parameters like lengths, k-mers and the alignment algorithm, changing the 'Target database' of the BLAST search can also point focus on fewer but more diverse sequences. Similarly, by using PSI-BLAST also evolutionary more distant homologs can be identified[1].

The maximum amount of sequences to retrieve from a single BLAST run from the webserver is currently limited to 5000 (NCBI) or 1000 (UniProt) entries. To conveniently study entire families or superfamilies, more distant evolutionary patterns, defining such a family, must be selected. Enzyme identification via motives has a long history: Databases including such a multitude of these definitions were PFAM (offline since 2023) and InterPro[5]. InterPro combines 13 protein signature databases into one central resource to facilitate definition and characterization of enzymes via typical patterns and signatures (e.g. active site residues or entire folds)[5]. The search and pool creation process can also be structurally motivated via structural homology searches using Foldseek[43] and ESM Metagenomic Atlas[26]. Foldseek cluster clusters datasets from a structural base like described before for sequences[4]. The Protein Universe Atlas[9] e.g. navigates such a clustered dataset at a diversity of 90% similarity (AFDB90) and is very handy to use.

#### Sequence comparison and visualization with SelectZyme

Once an enzyme pool has been assembled, the next practical bottleneck is to compare sequences at scale and translate these relationships into views that support human interpretation and selection. SelectZyme assists the comparison and interpretation step by representing all sequences as pLM embeddings and exposing their relationships through three linked views: a low-dimensional landscape (global context), an MST-based connectivity projection (local-to-global links), and a dendrogram (hierarchical organization). Selections and filters propagate across all views, enabling iterative refinement without re-running the full pipeline.

SelectZyme supports *sequence comparison* by allowing users to choose the underlying representation and similarity signal: sequence-only embeddings from pLMs such as ESM-1b, ESM2, or ProtT5, and, when desired, sequence-and-structure–aware representations via ProstT5. These embeddings are then translated into interpretable 2D coordinate systems using dimensionality-reduction methods (UMAP, t-SNE, or PCA), while local neighborhood structure is formalized through HDBSCAN, whose key controls (e.g., neighborhood size and minimum cluster density) tune the granularity of recovered groups and the separation between dense cores and outliers.

For *visualization*, SelectZyme provides three complementary representations that map the same relationships onto different, decision-relevant views. Graph-based views emphasize connectivity, nearest-neighbor links, and potential "walks" along related variants; clustering-based views highlight density structure, centroid representatives, and phylogeny-like distances via the dendrogram; and landscape views provide a global overview that can be explored by annotations to localize characterized regions and guide region-focused exploration.

#### Candidate selection strategies

SelectZyme is not limited to a single "best" mining route, but rather supports a spectrum of exploration and prioritization strategies for candidate selection that can be chosen depending on the experimental objective (e.g., broad novelty scouting, close-homolog optimisation, remote-homolog discovery, or IP-aware avoidance). Global diverse sampling, local homolog selection, and neighbourhood exploration of close or distant homologs can all be performed on the same landscape without re-running the pipeline or enforcing homology cut-offs, enabling rapid iteration between exploration and exploitation. Independent of the specific strategy, the joint use of density distributions (as a proxy for evolutionary sampling), community structure, cluster centroids, and connectivity information provides an interpretable basis to rank sub-regions and select representatives, while additional filters (organism, annotation status, characterised entries, etc.) allow tailoring the search toward practical constraints. Additionally, constraints can be imposed including both desired properties (e.g. if present: specific taxa, annotations, or functional neighborhoods) and "avoid" criteria. Below, we outline common mining scenarios:

***Global diverse sampling:*** Use a global view to sample distant regions of sequence space. Density distributions (reflecting where evolution provides many templates), cluster centroids, and network connectivity support the selection of informative representatives. Sampling complete outliers is possible, but often less informative because homology-based expansion from an isolated hit is limited.

***Local homolog selection:*** SelectZyme can capture structure-consistent similarity and may therefore yield different neighborhoods than BLAST, particularly in low sequence identity regime. In addition, relatedness is assessed not only relative to a single query (query vs. all) but across the entire dataset (all vs. all), facilitating interpretation of inter-search variance and neighborhood structure.

***Local close homolog exploration:*** Because SelectZyme does not hide sequences above a predefined identity threshold (as in clustered reference resources such as UniRef), dense local neighborhoods of close homologs can be explored alongside more distant homolog regions without changing hyperparameters or re-running the tool.

***"Avoidome":*** Incorporating patent-derived sequences or IP-relevant annotations as custom data can help identify underexplored regions for global sampling or flag local neighborhoods where remote homologs (divergent sequence, conserved fold) may be particularly interesting.

Across all strategies, density distributions, community structure, cluster centroids, and connectivity patterns help prioritise sub-regions, and the focus can be further refined using metadata filters (e.g., desired organisms, characterised entries, or specific annotations).

#### State-of-the-art tools useful for enzyme mining

Historically, identification of enzymes of interest relied on BLAST-based similarity searches[2] and multiple sequence alignments (MSAs), with phylogenetic trees commonly used for visualization via hierarchical clustering. As sequence databases expanded rapidly, these alignment-centric workflows became increasingly impractical at scale to cover protein families or superfamilies.

To address scalability, sequence-based clustering algorithms such as CD-HIT,[24,25] MAFFT, and MMseqs,[28] as well as traditional unsupervised machine learning algorithms like DBSCAN and HDBSCAN[13] have enabled large-scale redundancy reduction and the generation of clustered reference resources such as UniRef[42] and RefSeq.[8,31,33] However, clustering can obscure functional diversity by collapsing heterogeneous groups into a single representative sequence, which is often selected by sequence length rather than by representativeness across relevant biochemical features.[24,25]

Sequence similarity networks (SSNs) integrate alignment-based similarity measures with graph representations by modeling sequences as nodes connected through edges when pairwise similarity exceeds a defined threshold (e.g., BLAST e-value).[23] This framework has become a popular strategy for exploring functional and evolutionary relationships within large protein families, supported by dedicated tools such as EFI-EST[15,44] for automated network constructions and platforms like Cytoscape[37,40] for interactive visualization. However, as dataset size and diversity increase, SSNs rapidly become dense and visually cluttered, limiting interpretability and obscuring meaningful structure.[7] Although representative-node schemes and threshold tuning are commonly applied to mitigate graph complexity, the selection of robust and biologically meaningful parameters remains largely heuristic, particularly for large, heterogeneous families.[7] Hornung and Terrapon[23] clearly illustrates the extent to which manual optimization is often required to obtain interpretable networks. More fundamentally, because SSNs rely on pairwise sequence alignment, they inherit well-known limitations of alignment-based approaches, including sensitivity to variable sequence length, low sequence identity, and complex or modular family architectures.[10,11,17,22]

EnzymeMiner v1.0,[20] a dedicated mining tool for EC-number-based enzyme discovery, integrates sequence retrieval, additional property predictions and export formats for SNN visualization. Its computational workflow consists of PSI-BLAST[1], MSAs, clustering, and solubility prediction via SoluProt.[21] However, when no annotated example exists for a target EC number, it may require predefined essential residues, thereby presupposing substantial prior knowledge of the target. A recent v2.0 extends data retrival to metagenomic data and integrates additional protein property prediction tools.

Protein language models, including ESM2[26] and ProtT5[18], offer an attractive alternative by enabling alignment-free sequence representations through embeddings[34]. Tools such as ProteinClusterTools[7], Evolocity[19], and the interactive ProtSpace[36] demonstrate that embedding-based methods can capture latent structure in protein space and enable visualization via dimensionality reduction. While dimensionality reduction excels to indicate the presence of distinct groups in the protein family, it falls short to deal with the hierarchical structure of the sequence[41].

Yet, tools were designed primarily for general protein analysis and evolutionary modeling, rather than enzyme-specific mining tasks (except EnzymeMiner) that require richer interactivity, selection tracking and streamlined data export.

### SelectZyme implementation

SelectZyme was developed to bundle the presented workflow into an user-friendly application, addressing key limitations of current enzyme-mining workflows within a single, intuitive platform, including i) the visualization bottleneck that arises when similarity networks become too dense, ii) the loss of local diversity when large datasets are compressed into cluster representatives that may be chosen by arbitrary heuristics, and iii) the reliance on sequence alignments, which can be slow at scale and unreliable for low-homology relationships or highly variable sequence lengths. In doing so, SelectZyme intentionally bridges familiar visualizations that many experimentalists already use, such as phylogeny-inspired views, graph-based networks, and embedding-driven latent space projections of a sequence space.

Modular components for preprocessing, embedding extraction through pretrained protein language models, clustering, and visualization are bundled. In the follow sections we describe the main design objectives and key implementation aspects of these modules.

#### Input, data retrieval, and preprocessing

SelectZyme supports three input modes, including i) custom data upload, ii) automated sequence retrieval, and iii) a combination of both. Custom data can be provided as `.fasta` files or as tabular `.csv`/`.tsv` files. For tabular uploads, the minimal required fields are an identifier and the protein sequence, provided in columns named `accession` and `sequence`. Integration of proprietary sequences, user-curated annotations, or data from domain-specific resources and other sources beyond UniProt are enabled that way.

Alternatively, SelectZyme can retrieve sequences directly from UniProt using standard UniProt query terms. The query terms also enable filtering of data during retrieval, such as sequence length specifications e.g.. Users may additionally specify annotation fields to be returned according to the UniProt query-field convention. All retrieved and processed data can be exported either as `.fasta` or in tabular format (Excel) and re-imported as custom input to avoid repeated retrieval steps.

Preprocessing is applied automatically to prepare sequences for embedding extraction. However, it can be disabled. By default, SelectZyme removes sequences containing undetermined residues (X), as well as duplicate entries (duplicate accessions) and duplicate sequences. Optional filters allow removal of sequences lacking an N-terminal start methionine (M), which can simplify downstream gene synthesis and reduce manual curation efforts. For the default embedding model (ESM-1b), sequences longer than 1024 amino acids are excluded due to model constraints; when alternative pLMs are selected (ESM-2, ProtT5, or ProstT5), the effective sequence length limit is not fixed but primarily determined by available Graphics Processing Unit (GPU) memory. It is also possible to define a lower limit for sequence length to avoid too many peptidal artifacts during sequence retrieval.

#### Embedding via pre-trained protein language models

Protein sequences are represented as numerical embeddings using pretrained protein language models (pLMs). By default, SelectZyme uses ESM-1b[34], trained on UniRef50. Additional supported pLMs include ESM2[26], ProtT5[12], and ProstT5[18] (see Table **S1**). For all models, residue-level embeddings are aggregated by mean pooling over the full sequence to obtain a fixed-length per-sequence vector representation from the last layer of the pLM. For ESM-1b for instance, the residue embedding tensor of size $L \times 1280$ (with $L \leq 1024$) is reduced to a 1280-dimensional vector. For ProtT5 and ProstT5, SelectZyme additionally provides an option to compute the mean while excluding padding tokens. All embeddings are generated using the `transformers` library (v4.48.0) with the Hugging Face models detailed in Table **S1**.

**Table S1:** List of pre-trained models available in SelectZyme

| Model name | Huggingface version | Reference |
|---|---|---|
| ESM-1b | esm1b_t33_650M_UR50S | 34 |
| ESM2 | facebook/esm2_t33_650M_UR50D | 26 |
| ProtT5 | Rostlab/prot_t5_xl_uniref50 | 12 |
| ProstT5 | Rostlab/ProstT5 | 18 |

#### Orthogonal unsupervised machine learning methods

##### Dimensionality reduction

SelectZyme hosts three dimensionality-reduction strategies, including the linear method principal component analysis (PCA) and the nonlinear methods uniform manifold approximation and projection (UMAP) and t-distributed stochastic neighbor embedding (t-SNE). Default hyperparameter settings are the same for the methods and provided with: `random_state`=42 and `n_neighbors`=15 but can be adjusted by the user if desired. All dimensionality reductions are performed using GPU-accelerated implementations from cuML, where also a fallback on CPU is possible if no GPU is detected or available.

##### Density-based clustering

SelectZyme uses HDBSCAN clustering algorithm[27]. HDBSCAN extends DBSCAN[13] by formulating density-based clustering as a hierarchical problem and extracting a flat clustering based on cluster stability[27]. HDBSCAN has been applied successfully to biological sequence embeddings, including recent demonstrations on ESM-derived representations, and is suited for heterogeneous datasets that contain both dense neighborhoods and sparse regions[38]. We use the GPU-accelerated cuML implementation of HDBSCAN (v23.12), which requires the `hdbscan` Python package (v0.8.33). Default parameters for the clustering are provided in SelectZyme with `min_samples`=2 and `min_cluster_size`=2. For very large datasets it is advised to increase the values to make clustering appear more granular.

As part of the HDBSCAN library it is possible to compute a MST and perform Single-Linkage Clustering (SLC), which is used by SelectZyme to construct the dendrogram ('phylogeny' view). SLC has been described as a valid approach for phylogeny-inspired tree reconstruction from a distance matrix[3,30,35]. Conceptually, MST and SLC are closely related, as the single-linkage tree is contained within the corresponding MST[6,16]. Compared with classical phylogenetic methods such as neighbor joining or UPGMA, this approach offers computational advantages when large numbers of sequences are compared, since it leverages the same distance computations already required for clustering and connectivity estimation[16].

MSTs have recently been highlighted as effective visualization for large, high-dimensional datasets, including biological applications[32]. In SelectZyme, the MST is visualized using the mutual reachability distance, which differs from classical sequence-identity measures and instead reflects local connectivity in embedding space (i.e., how each sequence connects to its nearest neighbors under the clustering metric). Given its efficient computation and its ability to restore the global connectivity structure on top of low-dimensional projections, we incorporated MST-based visualizations. Related applications and insights of MST-based connectivity and potential 'graph walks' have been demonstrated previously[29].

#### Visualization and interactivity

Outputs from dimensionality reduction and clustering are rendered as scatter and line-based plots. Interactive visualizations are implemented with Plotly scatter plots using WebGL to enable client-side rendering of large datasets with GPU acceleration when available. To capture and persist user interactions, the plots are embedded in a Dash application with event listeners that record users´ point selections. Selected sequences are tracked in an editable table, enabling iterative curation and export during exploration.

#### Implementation, deployment, and reproducibility

SelectZyme is implemented in Python 3.10, with all dependencies documented in the public repository. Computationally intensive steps leverage NVIDIA RAPIDS GPU-accelerated libraries (CUDA 11.4–11.8), including cuDF (v23.12) and cuML (v23.12). Pretrained pLM embeddings are generated using Hugging Face models via the `transformers` library.

SelectZyme can be installed from source or deployed via Docker, with side packages managed through `conda` and `pip`. A GPU is recommended for practical runtimes on large datasets. For reproducibility, nondeterministic algorithms use a fixed random seed that can be specified in a YAML (`.yml`) configuration file, which also records parameter settings and reduces reliance on extensive command-line arguments. In addition to local usage instructions, a Jupyter notebook provides an overview of core computations and plots; however, full functionality—particularly interactive selection and tracking—requires running the SelectZyme web application.

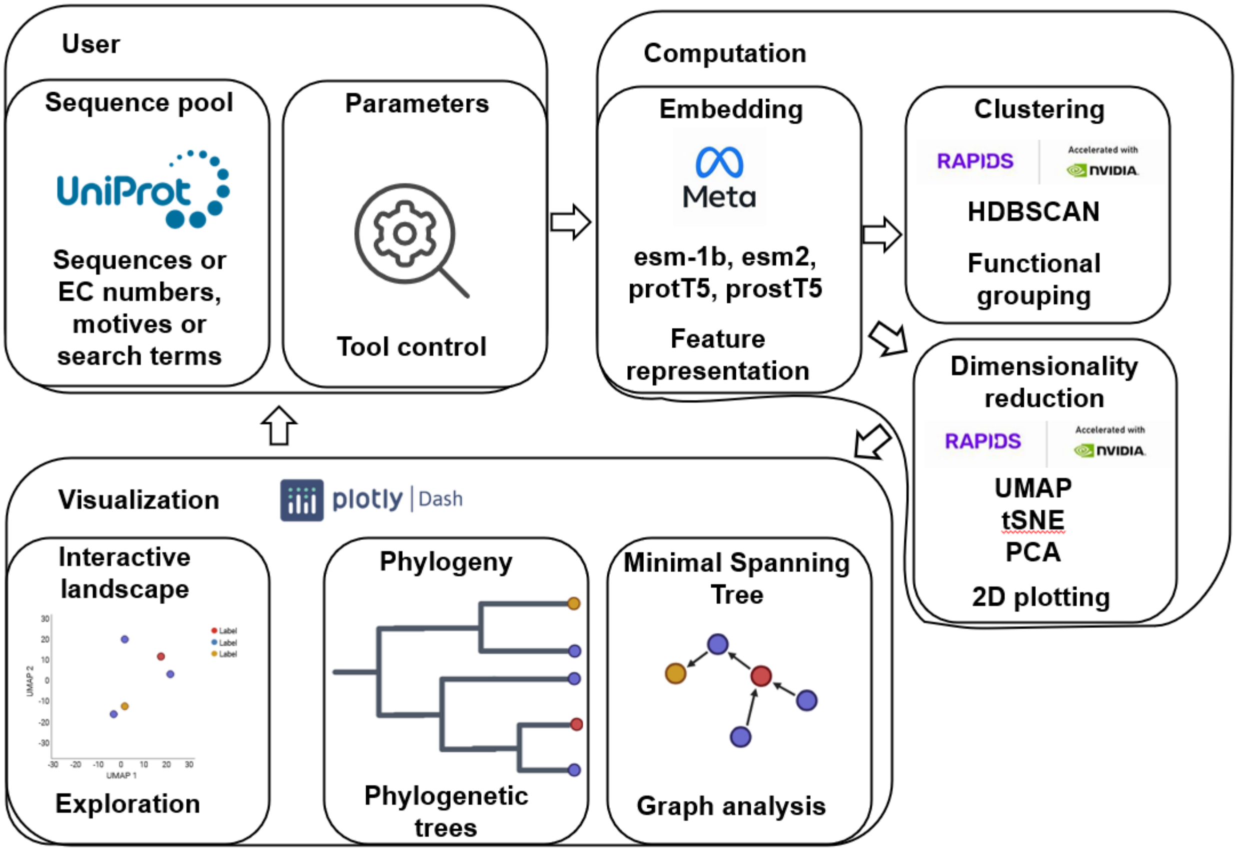

**Figure S2: Schematic workflow of computational steps in SelectZyme.**
A user interacts by defining a sequence pool and optionally adjusting parameters. Computational steps consist of embedding, clustering and dimensionality reduction. Visualization is performed in three different view modes: a 2D scatter plot, a phylogenetic tree and a connectivity graph, represented by a minimum spanning tree.

### Additional information about the selected sequences of the PETase dataset

Figure **S3** combines the example selection process of the PETase dataset from the main manuscript with a global overview A-C, and a refined view D-F for actual sequence selection. We selected three archaeal sequences as an example: two from Thermoproteota and one from Euryarchaeota (*Pyrococcus furiosus*). *P. furiosus* is well known for its thermostable DNA polymerase (Pfu), which supports Polymerase Chain Reaction (PCR) through enhanced thermostability and proofreading compared to Taq polymerase. With an optimal growth temperature of 100 °C, *P. furiosus* is classified as a hyperthermophile[14], suggesting that it encodes a broader repertoire of thermostable enzymes. Because it is also the closest neighbor to an experimentally PETase-active cluster centroid (X) (accession 387; Figure **S3** G), it represents an attractive candidate for experimental characterization.

The two Thermoproteota candidates were a *Thermofilum* sp. and *Pyrobaculum ferrireducens.* A strain of *P. ferrireducens* (1860) was isolated at 84 °C and pH 6.8 from sediment near a hot spring, and isolates were reported to grow optimally at 90–95 °C and pH 6.0–7.0[39]. Members of *Thermofilum* are also hyperthermophilic, with reported optima spanning high temperature and, in some cases, mildly acidic conditions: *Thermofilum adornatum* isolate 1910bT grows optimally at 80 °C and pH 5.5–6.0[46], *Thermofilum pendens* at 88 °C[47], and the hyperthermophilic crenarchaeon strain 3507LTT grows at 73–93 °C under pH 5.0–7.5 with an optimum at 85 °C and pH 6.0–6.7[45]. Tolerance to acidic conditions can be advantageous because PET depolymerization can acidify the reaction environment over time. All in all these small example selections should exemplyify aforementioned mining scenario of looking for extremophiles in a dataset, being close to manually curated PETase active regions, identified sequences have the potential to exhibit PETase activity, as well.

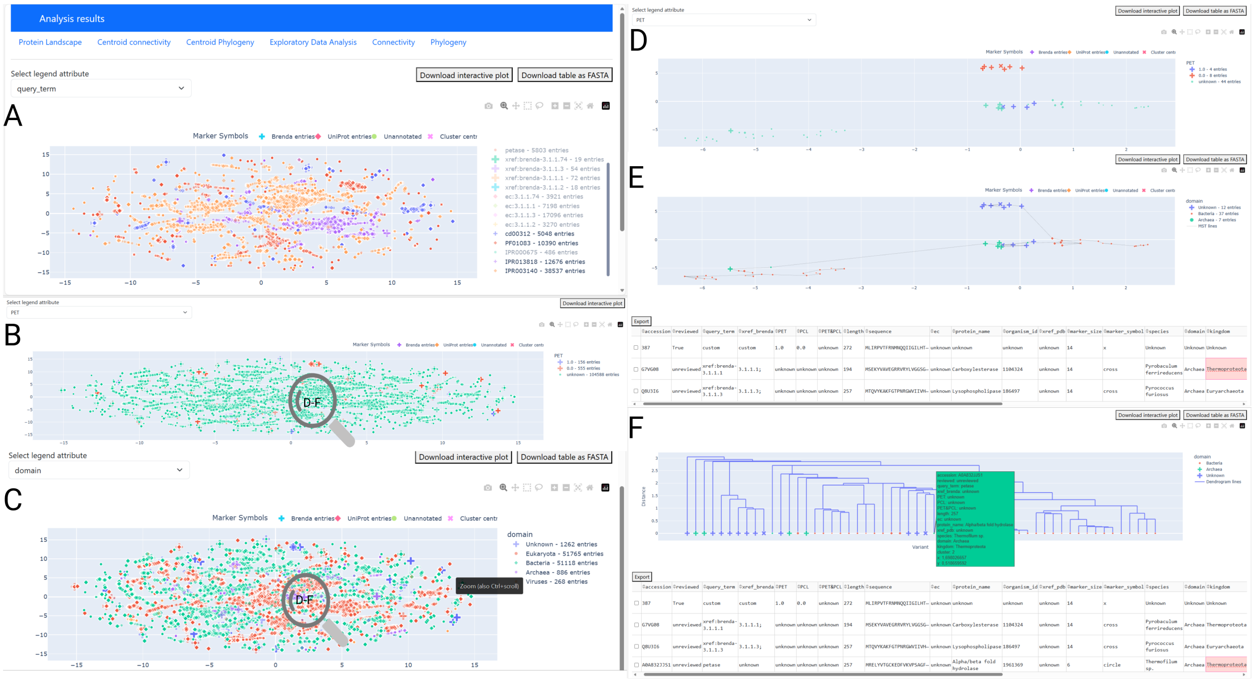

**Figure S3: SelectZyme: interactive navigation of large protein sequence spaces.** **A** Global view of the PETase dataset; the highlighted search term corresponds predominantly to InterPro motifs associated with cutinases (PF01083), esterases/lipases (cd00312, IPR013818), and carboxylesterases (IPR003140). **B** Global overview of PETase activity labels, where ‘1.0‘ indicates experimentally confirmed PETase activity, ‘0.0‘ indicates no activity, and ‘unknown‘ denotes sequences that have not been tested. **C** Global view highlighting the taxonomic ‘domain‘ to facilitate identification of archaeal candidates. **D** Magnified subregion showing co-occurrence of PETase activity, non-activity, and archaeal presence, obtained via a subsequent run on a box-selected region. **E** Minimum Spanning Tree projection onto the dimensionality-reduced space visualizes nearest-neighbor connectivity inferred from clustering; the sequence selector highlights three selected sequences (an experimentally active centroid (387) and two close neighbors, annotated as a archaeal sequences with an BRENDA-entry, too). **F** The dendrogram provides phylogeny-like distances for all sequences in the selected subregion; an additional archaeal candidate from the kingdom of Thermoproteota was selected based on proximity to the experimentally characterized cluster centroid.

### Additional representative applications of SelectZyme for other enzymes

Three additional datasets were visualized with SelectZyme to demonstrate diverse input sources, database query strategies, and subtype separability. The analyses are available for interactive exploration at the website. Overall, these additional datasets demonstrate that SelectZyme can accommodate very different input sources (free-text queries, EC/InterPro-guided retrieval, and domain-specific databases) while still yielding navigable landscapes for candidate selection across diverse mining strategies.

#### LPOR: Minimalistic query example

The Light-dependent Protochlorophyllide OxidoReductase (LPOR) dataset was included to demonstrate an enzyme family retrieved from minimal information sources, using only query terms such as "light-dependent protochlorophyllide oxidoreductase LPOR", "light-dependent protochlorophyllide oxidoreductase", and "lpor". No additional constraints (e.g., EC numbers or InterPro motifs) were specified, yielding 1,279 sequences of length 100–501 aa. SelectZyme default preprocessing was applied, removing duplicate UniProt IDs, duplicate sequences, sequences lacking a starting methionine (M), and sequences containing undetermined residues (X). Embeddings were generated with ESM2. Clustering was performed with HDBSCAN (`min_samples`=10, `min_cluster_size`=15, and dimensionality reduction with UMAP (`random_state`=42; `n_neighbors`=15). All additional settings are specified in `config.yml`.

#### FMO: Enzyme family subtype visualization

The Flavin-Containing Monooxygenase (FMO) dataset, in contrast, included InterPro motifs that distinguish FMO subtypes, and additional query terms and EC numbers were used to enrich the dataset to 66,086 sequences. FMOs illustrate subtype separability within a family, with the dominant head class spanning most of the space and accounting for the largest fraction of data points. FMO sequences and subtype-related entries (FMO) were retrieved from UniProt with a length filter of 200–601 aa using the following query terms: "ec:1.14.13.8", "IPR002254" (FMO2), "IPR002256" (FMO3/FMO4), "IPR002253" (FMO1), "IPR002257" (FMO5), "IPR000960", "Flavin-Containing Monooxygenase (FMO)", and "Flavin-Containing Monooxygenase". SelectZyme default preprocessing was applied (duplicate UniProt IDs/sequences removed; sequences lacking N-terminal M removed; sequences containing X removed). Embeddings were generated with ESM-1b. Clustering and dimensionality reduction were performed using HDBSCAN (`min_samples`=10, `min_cluster_size`=15) and UMAP (`random_state`=42; `n_neighbors`=15), respectively. The complete parameterization is documented in `config.yml`.

#### IRED: Comparison of different input sources (databases)

The Imine Reductase (IRED) dataset was used to demonstrate integration of a domain-specific database with InterPro motif annotation: BioCatNet IRED entries were visualized alongside InterPro motifs associated with IREDs, yielding 2,239 sequences. Interestingly, little overlap emerged, and interpretation remains challenging because the domain-specific source provides limited metadata. The key takeaway is that information sources matter, and third-party databases can substantially shape and enrich the mining dataset.

IRED sequences with a length of 200–701 aa were retrieved from UniProt using the queries "imine reductase" and "imine reductases", together with the corresponding motifs PF21390 and IPR048655. SelectZyme default preprocessing was applied (duplicate UniProt IDs/sequences removed; sequences lacking N-terminal M removed; sequences containing X removed). Embeddings were generated with ESM2. Clustering was performed with HDBSCAN (`min_samples`=10, `min_cluster_size`=15), and dimensionality reduction with UMAP (`random_state`=42; `n_neighbors`=15). Additional settings are provided in `config.yml`.

```
Reference sequence (1): 4
Identities normalised by aligned length.
Colored by: identity

                     cov    pid   1 [        .         .         .         .         :         .         .         . 80
1 4              100.0% 100.0%     --------------MLCLFLCGCVMKKWCGLAFSIFFSF--FLRAQDLTFQEVFEVAACNDD-VSLLNKMHKRTVRDNQF
2 5               96.8%  34.1%     -------------------MFERIKRYFLIIGFFCFFGLYVFTENKTDDFIGSYEIKAYNDVSRIGFNNVYKSIVKDNTY
3 6               91.3%  33.0%     --------------------------------------MNSHLQDEHKIFEGPYDIIAYNDKNIIGFNRVYKKTLEDNKE
4 7_outlier       84.6%   7.5%     MQTQHTKVTQTHFLLWILLLCMPFGKSHTEEDF---------------------IITT-------------------KTG
5 3               42.9%   6.6%     ------------------------------------------------------MSEK-------------------YVA
6 2               61.4%   7.5%     ------------------------------------------------------MLKQ-------------------SFQ
7 1               60.9%   8.7%     ------------------------------------------------------MKIK-------------------NQI
8 1a              61.2%   7.8%     ------------------------------------------------------MIKK-------------------NQI
  consensus/100%                   .........................................................t......................
  consensus/90%                    .........................................................t......................
  consensus/80%                    .......................................................hht...................p..
  consensus/70%                    .......................................................ltt...................sph

                     cov    pid  81          .         1         .         .         .         .         :         . 160
1 4              100.0% 100.0%     DEQLTDRVHLRYPASVYYPAHQGTYPLIML-------------------------------------SHGWN--------
2 5               96.8%  34.1%     DKTLSDKENLRGEITVYLPKKTGKYPLIMI-------------------------------------THGWA--------
3 6               91.3%  33.0%     DKSLSDEENLRDEVTVYIPKKKGKYPLILV-------------------------------------SHGWA--------
4 7_outlier       84.6%   7.5%     RVRGLSMPVLGGTVTAFLGIPYAQPPLGSLRFKKPQPLNKWPDIHNATQYANSCYQNIDQAFPGFQGSEMWNPNTNLSED
5 3               42.9%   6.6%     ---------VEGRRVRYLVGGSGRP-VVL------------------------LH-GWSFNADTWVESGVFN--------
6 2               61.4%   7.5%     MDVGKEERVIRGEVTIPSNLSNPVPTIII------------------------CH-GFK----AFKEWGFFP--------
7 1               60.9%   8.7%     LTRINQKPIVY--DTFYNDNKKQKPLVIF------------------------CH-GYK----GFKDWGTWD--------
8 1a              61.2%   7.8%     LNRNNQKPIVF--DTFFNKTEKQKPLVVF------------------------CH-GYK----GFKDWGCWN--------
  consensus/100%                   .........l....sh..........lh........................................thas........
  consensus/90%                    .........l....sh..........lh........................................thas........
  consensus/80%                    .....p...lh...oha.shtptp..llh.......................................thas........
  consensus/70%                    .pphppc..lt..sTha.stppsp..llh.......................................tsWs........

                     cov    pid 161          .         .         .         2         .         .         .         . 240
1 4              100.0% 100.0%     -------------------------STQDYQ----RALARFLAAQGFVTVVFTS---RAQRRPPDFLS--TFDSV-----
2 5               96.8%  34.1%     -------------------------NSKLAF----LSLGRYIASHGYAAAVFTS---KKRSLPKDWLP--AFSSV-----
3 6               91.3%  33.0%     -------------------------NYKYGL----LPIGHYLASYGYAVALFTS---KKKAIPKDWIP--TFTAV-----
4 7_outlier       84.6%   7.5%     CLYLNVWIPVPKPKNATVMVWIYGGGFQTGTSSLPVYDGKFLARVE--RVIVVSMNYRVGAL--GFLAFPGNPDAPGNMG
5 3               42.9%   6.6%     -----------------------------------SLAGHFAV-------YAIDMPYGIRTKSERFQA-----------P
6 2               61.4%   7.5%     -----------------------------------ILAEKLADAGFVAITFDFSMNG-VGDDPNTYSELEKFARLTFSRE
7 1               60.9%   8.7%     -----------------------------------LVANNFAKQNLFFVKFNFSHNGGTANQPIDFPDLDAFAENNYTKE
8 1a              61.2%   7.8%     -----------------------------------LMAEAFMNANLFFVKFNFSHNGGTIENPIDFPDLNAFAENNYTKE
  consensus/100%                   ......................................tthh........h.hs....h.t...ta..............
  consensus/90%                    ......................................tthh........h.hs....h.t...ta..............
  consensus/80%                    ...................................h.htpahst....hhh.hS...thtt.s.sa.s..t.st......
  consensus/70%                    ...................................lhhs+ahst.hhhhhh.hS...thts.P.ca.s..sFsps.....

                     cov    pid 241          :         .         .         .         .         3         .         . 320
1 4              100.0% 100.0%     ---YALMQRVNEQEGSPLYRKVDFTRVGVLGHSMGGTAALHYANRYPERIRTV--------------VALHPFNGG----
2 5               96.8%  34.1%     ---YGIINEAVCNKNHSMYNSIDMKNIGIIAHSMGGAAALYYANFIPE-VKAV--------------AAIHPYNGS----
3 6               91.3%  33.0%     ---HNLIKNKNEDASHDLYGLIDINNIGIVTHSMSGPASFYYASLRPE-VKAI--------------SAIHPYNGA----
4 7_outlier       84.6%   7.5%     LFDQQLALQWVQRNIAAFG--GNPKSITIFGESAGAASVSLHLLC-PQSYPLFTRAILESGSSNAPWAVKHPEEARNRTL
5 3               42.9%   6.6%     RREYAVFLRRV-------LDALDLADPPLVGPSASGEVVLWYVAKRLPTRAA------------VVVGPV--------GL
6 2               61.4%   7.5%     QEDIALILQAIQQGQLPYSSQMDKERLGILGHSRGGGNSLIFTLEHPQIKSV------------TIWNSIPRPDFF----
7 1               60.9%   8.7%     LQDLDDILNHFTSDQSIYKNNIDTNNITLIGHSRGGGISIIKASEDHRITKL------------ITWASVCAFGKR--TS
8 1a              61.2%   7.8%     LDDLDAILNYFLANDYEYKNEININNIVVIGHSRGGGIAIIKSAEDARIKKL------------ITFASVSDFSKR--TS
  consensus/100%                   ....t.h.ph...........hs.tp.slhs.Shuu..s.hh.......h.h...............ssh..........
  consensus/90%                    ....t.h.ph...........hs.tp.slhs.Shuu..s.hh.......h.h...............ssh..........
  consensus/80%                    ...hshh.ph..tt...hht.hshpplsllucShuGs.shhhs....p.hth...............ssl...th.....
  consensus/70%                    ...hslhhphhpptt..hhstlDhpslsllGHShGGssulhassp.sp.hph...............uulps.stt....

                     cov    pid 321          .         .         :         .         .         .         .         4 400
1 4              100.0% 100.0%     --------ADMVNMVGGKNEQLGTDLRSVRAAVLILTGT--KDMLAYPERSYEFFQSLPQDVPACFLSLKGMGHGA----
2 5               96.8%  34.1%     --------SVLVETVGSSNEELGDSFTETNAAVLILTSE--IDITAYPEKTYRFFKNLNKNNPACFLSFQNVKHNG----
3 6               91.3%  33.0%     --------SSFVEAIASSNEELGDEFPKIKSAVLFLTSE--IDRSAYPEKTYKFFKNLNKDAPACFLSFKNIKHNG----
4 7_outlier       84.6%   7.5%     TLAKFTGCSKENEMEMIKCLRSKDPQEILRNERFVLPSDSILSINFGPTVDGDFL----TDMPHTLLQLGKVKK------
5 3               42.9%   6.6%     AGELLE-ELRGAGVPILAVWGEGDE----------------------------------VSPPSRASLLRDVA----KVV
6 2               61.4%   7.5%     GDELIR-EIKEKGRVYITNARTKQ------------------------------------EMPIDLEVIEDLERNWERFN
7 1               60.9%   8.7%     TTGDLE-QWKQDGVKYVLNSRTKQ------------------------------------NMPHNFQFYLDYINNKDRLD
8 1a              61.2%   7.8%     TIGDLE-EWKKLGVKYVLNGRTKQ------------------------------------QMPHNYQFYKDFKANEERLN
  consensus/100%                   ............th.h..s.t.tp....................................p.P.ph..h.th........
  consensus/90%                    ............th.h..s.t.tp....................................p.P.ph..h.th........
  consensus/80%                    .........p..th.h..s.p.tp....................................p.Ptsh..htshtt......
  consensus/70%                    .........c.sthhhhtNtchtp....................................shPtsh..hpshtps.....
```

#### Figure S4: Multiple Sequence Alignment of selected sequences 1-7 given in Figure 3

Complete Clustal Omega MSA of the 8 selected sequences (Fig. 3) from the selected sub-region of the PETase dataset. Shown are the alignable residues along the entire sequence length.

```
                   cov    pid 401          .         .         .         .         :         .         .         . 480
1 4             100.0% 100.0%     -----------------GTDK------YFEVWGFTYNKT-LSVESFALYRRLISAWFR---LF-LREEEEYRALFNE-HG
2 5              96.8%  34.1%     -----------------SLDI------YRTPLSGG-----YNEPFFKLYAGLDLAWFD---VF-LKNKRDGLEYFKP-DG
3 6              91.3%  33.0%     -----------------ALDI------FKTPLSGG-----YDEKAFKLYSRLSVLWFD---AF-LKGKKENLKYFQI-DD
4 7_outlier       84.6%   7.5%     ----------AQILVGVNKDEGTAFLVYGAPGFSKDNDSLITRKEFQEGLNMYFPGVSR-----L-GKEAVLFYYVDWLG
5 3              42.9%   6.6%     IL--------RDAGHAAYLDK---------PGEFV-------------EVVLDFL-------------------------
6 2              61.4%   7.5%     FIKRLSTLTQPLLIVQADQDH---------PGFME-------------GAQQIFQQAPRARLEVISNANHTFRAVHPFTS
7 1              60.9%   8.7%     IENCCKRLTIPHLIIHGTKDP---------SVSFE-------------EAKLLKLWSPKNTLIALPETNHVFNAKHPWTE
8 1a             61.2%   7.8%     IEKSEKSLQIKHLIIHAKNDK---------SIKYN-------------EAEMLHNWSKHSELFPIENSDHVFSTKHPWLN
  consensus/100%                  ...................D..........sh................................................
  consensus/90%                   ...................D..........sh................................................
  consensus/80%                   .................t.D..........sh.ht..............hth.h..........l.ttpt.h.hh....t
  consensus/70%                   .................shDh.........shtht..............sph.h.h.....h..l.tppcshthhp...s

                   cov    pid 481          .         5         .         .         .         .         :         . 560
1 4             100.0% 100.0%     ----RPFRNLKPHL-----------------------------------------------------SQ-----------
2 5              96.8%  34.1%     ----EKFKEIKNFL-----------------------------------------------------YE-----------
3 6              91.3%  33.0%     ----EKFQDIKDLL-----------------------------------------------------YT-----------
4 7_outlier       84.6%   7.5%     EQSPEVYRDALDDVIGDYNIICPALEFTKKFAELENNAFFYFFEHRSSKLPWPEWMGVMHGYEIEFVFGLPLGRRVNYTR
5 3              42.9%   6.6%     -------------------------------------------------------------------RGVY---------
6 2              61.4%   7.5%     MT--PQLNDAINL-------------------------------------------------SVQFFQETLTG-------
7 1              60.9%   8.7%     NYLPEALEVVTKK-------------------------------------------------SIAFIN------------
8 1a             61.2%   7.8%     STMPKELKQVVDK-------------------------------------------------SINFINN-----------
  consensus/100%                  ................................................................................
  consensus/90%                   ................................................................................
  consensus/80%                   ......hp.hh.........................................................t...........
  consensus/70%                   ....c.hpphhsh.......................................................t...........

                   cov    pid 561          .         .         .         6         .         .         .         . 640
1 4             100.0% 100.0%     -----------------------ETGRA-HSYPFFCEQNISN--------------------------------------
2 5              96.8%  34.1%     -----------------------KPRGEHEVYPNYDSRNLE---------------------------------------
3 6              91.3%  33.0%     -----------------------EIHREHEAYPSYDSRNLK---------------------------------------
4 7_outlier       84.6%   7.5%     AEEIFSRSIMKTWANFAKYGHPNGTQGNSTMWPVFTSTEQKYLTLNTEKSKIYSKLRAPQCQFWRLFFPKVLEMTGDIDE
5 3              42.9%   6.6%     --------------------------------------------------------------------------------
6 2              61.4%   7.5%     --------------------------------------------------------------------------------
7 1              60.9%   8.7%     --------------------------------------------------------------------------------
8 1a             61.2%   7.8%     --------------------------------------------------------------------------------
  consensus/100%                  ................................................................................
  consensus/90%                   ................................................................................
  consensus/80%                   ................................................................................
  consensus/70%                   ................................................................................

                   cov    pid 641          :         .         .     ] 676
1 4             100.0% 100.0%     ------------------------------------
2 5              96.8%  34.1%     ------------------------------------
3 6              91.3%  33.0%     ------------------------------------
4 7_outlier       84.6%   7.5%     TEQEWKAGFHRWSNYMMDWQNQFNDYTSKKESCTAL
5 3              42.9%   6.6%     ------------------------------------
6 2              61.4%   7.5%     ------------------------------------
7 1              60.9%   8.7%     ------------------------------------
8 1a             61.2%   7.8%     ------------------------------------
  consensus/100%                  ....................................
  consensus/90%                   ....................................
  consensus/80%                   ....................................
  consensus/70%                   ....................................
```

MView 1.67, Copyright © 1997-2020 Nigel P. Brown

**Figure S5: Multiple Sequence Alignment of selected sequences 1-7 given in Figure 3 (continued)**

Complete Clustal Omega MSA of the 8 selected sequences (Fig. 3) from the selected sub-region of the PETase dataset. Shown are the alignable residues along the entire sequence length.

### References

(1) Altschul, S. (1997). Gapped blast and psi-blast: a new generation of protein database search programs. *Nucleic Acids Research*, 25(17):3389–3402.

(2) Altschul, S. F., Gish, W., Miller, W., Myers, E. W., and Lipman, D. J. (1990). Basic local alignment search tool. *Journal of Molecular Biology*, 215(3):403–410.

(3) Antonelli, A., Hettling, H., Condamine, F. L., Vos, K., Nilsson, R. H., Sanderson, M. J., Sauquet, H., Scharn, R., Silvestro, D., Töpel, M., Bacon, C. D., Oxelman, B., and Vos, R. A. (2016). Toward a self-updating platform for estimating rates of speciation and migration, ages, and relationships of taxa. *Systematic Biology*, page syw066.

(4) Barrio-Hernandez, I., Yeo, J., Jänes, J., Mirdita, M., Gilchrist, C. L. M., Wein, T., Varadi, M., Velankar, S., Beltrao, P., and Steinegger, M. (2023). Clustering-predicted structures at the scale of the known protein universe. *Nature*, pages 637–645.

(5) Blum, M., Chang, H.-Y., Chuguransky, S., Grego, T., Kandasaamy, S., Mitchell, A., Nuka, G., Paysan-Lafosse, T., Qureshi, M., Raj, S., Richardson, L., Salazar, G. A., Williams, L., Bork, P., Bridge, A., Gough, J., Haft, D. H., Letunic, I., Marchler-Bauer, A., Mi, H., Natale, D. A., Necci, M., Orengo, C. A., Pandurangan, A. P., Rivoire, C., Sigrist, C. J. A., Sillitoe, I., Thanki, N., Thomas, P. D., Tosatto, S. C. E., Wu, C. H., Bateman, A., and Finn, R. D. (2020). The InterPro protein families and domains database: 20 years on. *Nucleic Acids Research*, 49(D1):D344–D354.

(6) Chaudhuri, K., Dasgupta, S., Kpotufe, S., and von Luxburg, U. (2014). Consistent procedures for cluster tree estimation and pruning. *IEEE Transactions on Information Theory*, 60(12):7900–7912.

(7) Chen, J. Z., Gall, B., Pulsford, S. B., Tokuriki, N., and Jackson, C. J. (2025). Exploring large protein sequence space through homology- and representation-based hierarchical clustering. *Molecular Biology and Evolution*, 42(6):msaf136.

(8) Chen, Q., Wan, Y., Lei, Y., Zobel, J., and Verspoor, K. (2016). Evaluation of cd-hit for constructing non-redundant databases. In *2016 IEEE International Conference on Bioinformatics and Biomedicine (BIBM)*, pages 703–706. IEEE.

(9) Durairaj, J., Waterhouse, A. M., Mets, T., Brodiazhenko, T., Abdullah, M., Studer, G., Tauriello, G., Akdel, M., Andreeva, A., Bateman, A., Tenson, T., Hauryliuk, V., Schwede, T., and Pereira, J. (2023). Uncovering new families and folds in the natural protein universe. *Nature*, 622(7983):646–653.

(10) Edgar, R. C. (2010). Search and clustering orders of magnitude faster than BLAST. *Bioinformatics*, 26(19):2460–2461.

(11) Edgar, R. C. and Batzoglou, S. (2006). Multiple sequence alignment. *Current Opinion in Structural Biology*, 16(3):368–373.

(12) Elnaggar, A., Heinzinger, M., Dallago, C., Rehawi, G., Wang, Y., Jones, L., Gibbs, T., Feher, T., Angerer, C., Steinegger, M., Bhowmik, D., and Rost, B. (2022). Prottrans: Toward understanding the language of life through self-supervised learning. *IEEE Transactions on Pattern Analysis and Machine Intelligence*, 44(10):7112–7127.

(13) Ester, M., Kriegel, H., Sander, J., and Xu, X. (1996). A density-based algorithm for discovering clusters in large spatial databases with noise. *KDD*, pages 226–231.

(14) Fiala, G. and Stetter, K. O. (1986). Pyrococcus furiosus sp. nov. represents a novel genus of marine heterotrophic archaebacteria growing optimally at 100°c. *Archives of Microbiology*, 145(1):56–61.

(15) Gerlt, J. A., Bouvier, J. T., Davidson, D. B., Imker, H. J., Sadkhin, B., Slater, D. R., and Whalen, K. L. (2015). Enzyme function initiative-enzyme similarity tool (efi-est): A web tool for generating protein sequence similarity networks. *Biochimica et Biophysica Acta (BBA) - Proteins and Proteomics*, 1854(8):1019–1037.

(16) Gower, J. C. and Ross, G. J. S. (1969). Minimum spanning trees and single linkage cluster analysis. *Applied Statistics*, 18(1):54.

(17) Hamamsy, T., Morton, J. T., Blackwell, R., Berenberg, D., Carriero, N., Gligorijevic, V., Strauss, C. E. M., Leman, J. K., Cho, K., and Bonneau, R. (2023). Protein remote homology detection and structural alignment using deep learning. *Nature Biotechnology*, 42(6):975–985.

(18) Heinzinger, M., Weissenow, K., Sanchez, J. G., Henkel, A., Mirdita, M., Steinegger, M., and Rost, B. (2024). Bilingual language model for protein sequence and structure. *NAR Genomics and Bioinformatics*, 6(4):lqae150.

(19) Hie, B. L., Yang, K. K., and Kim, P. S. (2022). Evolutionary velocity with protein language models predicts evolutionary dynamics of diverse proteins. *Cell Systems*, 13(4):274–285.e6.

(20) Hon, J., Borko, S., Stourac, J., Prokop, Z., Zendulka, J., Bednar, D., Martinek, T., and Damborsky, J. (2020). EnzymeMiner: automated mining of soluble enzymes with diverse structures, catalytic properties and stabilities. *Nucleic Acids Research*, 48(W1):W104–W109.

(21) Hon, J., Marusiak, M., Martinek, T., Kunka, A., Zendulka, J., Bednar, D., and Damborsky, J. (2021). Soluprot: prediction of soluble protein expression in escherichia coli. *Bioinformatics*, 37(1):23–28.

(22) Hong, L., Hu, Z., Sun, S., Tang, X., Wang, J., Tan, Q., Zheng, L., Wang, S., Xu, S., King, I., Gerstein, M., and Li, Y. (2024). Fast, sensitive detection of protein homologs using deep dense retrieval. *Nature Biotechnology*, pages 983–995.

(23) Hornung, B. V. H. and Terrapon, N. (2023). An objective criterion to evaluate sequence-similarity networks helps in dividing the protein family sequence space. *PLOS Computational Biology*, 19(8):e1010881.

(24) Huang, Y., Niu, B., Gao, Y., Fu, L., and Li, W. (2010). CD-HIT suite: a web server for clustering and comparing biological sequences. *Bioinformatics*, 26(5):680–682.

(25) Li, W. and Godzik, A. (2006). Cd-hit: a fast program for clustering and comparing large sets of protein or nucleotide sequences. *Bioinformatics*, 22(13):1658–1659.

(26) Lin, Z., Akin, H., Rao, R., Hie, B., Zhu, Z., Lu, W., Smetanin, N., Verkuil, R., Kabeli, O., Shmueli, Y., dos Santos Costa, A., Fazel-Zarandi, M., Sercu, T., Candido, S., and Rives, A. (2023). Evolutionary-scale prediction of atomic-level protein structure with a language model. *Science*, 379(6637):1123–1130.

(27) McInnes, L., Healy, J., and Astels, S. (2017). hdbscan: Hierarchical density based clustering. *The Journal of Open Source Software*, 2(11):205.

(28) Mirdita, M., Steinegger, M., and Söding, J. (2019). Mmseqs2 desktop and local web server app for fast, interactive sequence searches. *Bioinformatics*, 35(16):2856–2858.

(29) Muir, D. F., Asper, G. P. R., Notin, P., Posner, J. A., Marks, D. S., Keiser, M. J., and Pinney, M. M. (2025). Evolutionary-scale enzymology enables exploration of a rugged catalytic landscape. *Science*, 388(6752):eadu1058.

(30) Nasıbov, E. and Kandemır-Cavas, C. (2011). Owa-based linkage method in hierarchical clustering: Application on phylogenetic trees. *Expert Systems with Applications*, 38(10):12684–12690.

(31) O'Leary, N. A., Wright, M. W., Brister, J. R., Ciufo, S., Haddad, D., McVeigh, R., Rajput, B., Robbertse, B., Smith-White, B., Ako-Adjei, D., Astashyn, A., Badretdin, A., Bao, Y., Blinkova, O., Brover, V., Chetvernin, V., Choi, J., Cox, E., Ermolaeva, O., Farrell, C. M., Goldfarb, T., Gupta, T., Haft, D., Hatcher, E., Hlavina, W., Joardar, V. S., Kodali, V. K., Li, W., Maglott, D., Masterson, P., McGarvey, K. M., Murphy, M. R., O'Neill, K., Pujar, S., Rangwala, S. H., Rausch, D., Riddick, L. D., Schoch, C., Shkeda,

A., Storz, S. S., Sun, H., Thibaud-Nissen, F., Tolstoy, I., Tully, R. E., Vatsan, A. R., Wallin, C., Webb, D., Wu, W., Landrum, M. J., Kimchi, A., Tatusova, T., DiCuccio, M., Kitts, P., Murphy, T. D., and Pruitt, K. D. (2015). Reference sequence (RefSeq) database at NCBI: current status, taxonomic expansion, and functional annotation. *Nucleic Acids Research*, 44(D1):D733–D745.

(32) Probst, D. and Reymond, J.-L. (2020). Visualization of very large high-dimensional data sets as minimum spanning trees. *Journal of Cheminformatics*, 12(1):12.

(33) Pruitt, K. D. (2004). Ncbi reference sequence (refseq): a curated non-redundant sequence database of genomes, transcripts and proteins. *Nucleic Acids Research*, 33(Database issue):D501–D504.

(34) Rives, A., Meier, J., Sercu, T., Goyal, S., Lin, Z., Liu, J., Guo, D., Ott, M., Zitnick, C. L., Ma, J., and Fergus, R. (2021). Biological structure and function emerge from scaling unsupervised learning to 250 million protein sequences. *Proceedings of the National Academy of Sciences*, 118(15):e2016239118.

(35) Sanderson, M. J., Boss, D., Chen, D., Cranston, K. A., and Wehe, A. (2008). The phylota browser: Processing genbank for molecular phylogenetics research. *Systematic Biology*, 57(3):335–346.

(36) Senoner, T., Olenyi, T., Heinzinger, M., Spannagl, A., Bouras, G., Rost, B., and Koludarov, I. (2025). Protspace: A tool for visualizing protein space. *Journal of Molecular Biology*, 437(15):168940.

(37) Shannon, P., Markiel, A., Ozier, O., Baliga, N. S., Wang, J. T., Ramage, D., Amin, N., Schwikowski, B., and Ideker, T. (2003). Cytoscape: A software environment for integrated models of biomolecular interaction networks. *Genome Research*, 13(11):2498–2504.

(38) Simon, E. and Zou, J. (2025). Interplm: discovering interpretable features in protein language models via sparse autoencoders. *Nature Methods*, 22(10):2107–2117.

(39) Slobodkina, G. B., Lebedinsky, A. V., Chernyh, N. A., Bonch-Osmolovskaya, E. A., and Slobodkin, A. I. (2015). Pyrobaculum ferrireducens sp. nov., a hyperthermophilic fe(iii)-, selenate- and arsenate-reducing crenarchaeon isolated from a hot spring. *International Journal of Systematic and Evolutionary Microbiology*, 65(Pt3):851–856.

(40) Su, G., Morris, J. H., Demchak, B., and Bader, G. D. (2014). Biological network exploration with cytoscape 3. *Current Protocols in Bioinformatics*, 47(1):8–13.

(41) Susmelj, A. K., Ren, Y., Vander Meersche, Y., Gelly, J.-C., and Galochkina, T. (2023). Poincaré maps for visualization of large protein families. *Briefings in Bioinformatics*, 24(3):bbad103.

(42) Suzek, B. E., Huang, H., McGarvey, P., Mazumder, R., and Wu, C. H. (2007). Uniref: comprehensive and non-redundant uniprot reference clusters. *Bioinformatics*, 23(10):1282–1288.

(43) van Kempen, M., Kim, S. S., Tumescheit, C., Mirdita, M., Lee, J., Gilchrist, C. L. M., Söding, J., and Steinegger, M. (2023). Fast and accurate protein structure search with foldseek. *Nature Biotechnology*, 42(2):243–246.

(44) Zallot, R., Oberg, N., and Gerlt, J. A. (2019). The EFI web resource for genomic enzymology tools: Leveraging protein, genome, and metagenome databases to discover novel enzymes and metabolic pathways. *Biochemistry*, 58(41):4169–4182.

(45) Zayulina, K. S., Elcheninov, A. G., Toshchakov, S. V., Kochetkova, T. V., Novikov, A. A., Blamey, J. M., and Kublanov, I. V. (2021). Novel hyperthermophilic crenarchaeon infirmifilum lucidum gen. nov. sp. nov., reclassification of thermofilum uzonense as infirmifilum uzonense comb. nov. and assignment of the family thermofilaceae to the order thermofilales ord. nov. *Systematic and Applied Microbiology*, 44(4):126230.

(46) Zayulina, K. S., Kochetkova, T. V., Piunova, U. E., Ziganshin, R. H., Podosokorskaya, O. A., and Kublanov, I. V. (2020). Novel hyperthermophilic crenarchaeon thermofilum

adornatum sp. nov. uses gh1, gh3, and two novel glycosidases for cellulose hydrolysis. *Frontiers in Microbiology*, 10:2972.

(47) Zillig, W., Gierl, A., Schreiber, G., Wunderl, S., Janekovic, D., Stetter, K., and Klenk, H. (1983). The archaebacterium thermofilum pendens represents, a novel genus of the thermophilic, anaerobic sulfur respiring thermoproteales. *Systematic and Applied Microbiology*, 4(1):79–87.
